# Enhancement of mediodorsal thalamus rescues aberrant belief dynamics in a mouse model with schizophrenia-associated mutation

**DOI:** 10.1101/2024.01.08.574745

**Authors:** Tingting Zhou, Yi-Yun Ho, Ray X. Lee, Amanda B. Fath, Kathleen He, Jonathan Scott, Navdeep Bajwa, Nolan D. Hartley, Jonathan Wilde, Xian Gao, Cui Li, Evan Hong, Matthew R. Nassar, Ralf D. Wimmer, Tarjinder Singh, Michael M. Halassa, Guoping Feng

## Abstract

Optimizing behavioral strategy requires belief updating based on new evidence, a process that engages higher cognition. In schizophrenia, aberrant belief dynamics may lead to psychosis, but the mechanisms underlying this process are unknown, in part, due to lack of appropriate animal models and behavior readouts. Here, we address this challenge by taking two synergistic approaches. First, we generate a mouse model bearing patient-derived point mutation in Grin2a (*Grin2a^Y700X+/−^*), a gene that confers high-risk for schizophrenia and recently identified by large-scale exome sequencing. Second, we develop a computationally trackable foraging task, in which mice form and update belief-driven strategies in a dynamic environment. We found that *Grin2a^Y700X+/−^* mice perform less optimally than their wild-type (WT) littermates, showing unstable behavioral states and a slower belief update rate. Using functional ultrasound imaging, we identified the mediodorsal (MD) thalamus as hypofunctional in *Grin2a^Y700X+/−^* mice, and *in vivo* task recordings showed that MD neurons encoded dynamic values and behavioral states in WT mice. Optogenetic inhibition of MD neurons in WT mice phenocopied *Grin2a^Y700X+/−^* mice, and enhancing MD activity rescued task deficits in Grin2a*^Y700X+/−^* mice. Together, our study identifies the MD thalamus as a key node for schizophrenia-relevant cognitive dysfunction, and a potential target for future therapeutics.

## Main

Genetics constitute more than 70% of the risk for schizophrenia^1^. Large impact genetic variants are rare in schizophrenia but key to generating animal models and studying neurobiological mechanisms. A recent large-scale exome sequencing study sampling ∼25,000 people with schizophrenia and ∼100,000 controls by the Schizophrenia Exome Sequencing Meta-analysis (SCHEMA) consortium identified 10 genes with rare coding variants that significantly increased risks for schizohrenia^2^. One of these genes is Grin2a with its coding variants associated with >20-fold increase in schizophrenia risks. Grin2a encodes glutamate postsynaptic N-methyl-D-aspartate receptor (NMDAR) subunit NR2A, which plays an important role in synaptic plasticity. NR2A is expressed after birth, and its levels continue to rise during postnatal development in the cortex and hippocampus^3^, a pattern consistent with the clinical course of schizophrenia which is featured by an asymptomatic state in childhood to full expression of the disease in late adolescence/early adulthood^4^. One of the point mutations Y700X, a stop codon-gaining protein-truncating variant, is located at the agonist binding domain of NR2A, likely resulting in NMDAR hypofunction, as would be consistent with one of the major theories of schizophrenia etiology —the glutamate hypothesis^5^. Using the CRISPR/Cas9 method, we generated a mouse model bearing the same point mutation (*Grin2a^Y700X^*), enabling us to model a substantial schizophrenia risk factor in mice.

Although positive symptoms of schizophrenia are a major focus of clinical diagnosis and treatment, cognitive impairment may occur earlier ^6,7^ and may even be an underlying cause of psychotic symptoms ^6–8^. Current antipsychotics target positive symptoms, but fail to treat cognitive dysfunction in most schizophrenia patients and 40% of patients are completely treatment resistant ^9^. Therefore, a better understanding of the mechanisms underlying cognitive deficits is key to developing novel therapeutic approaches ^7,10–12^.

One of the major roles of cognition is to generate flexible behaviors. Doing so requires integrating prior beliefs with new evidence to generate updated beliefs, a process whose dysfunction is thought to underlie delusional ideation ^13–15^. In fact, numerous studies have shown impaired belief updating in schizophrenia^13–17^, but the underlying neural mechanisms remain unknown. In order to tackle this challenge, we modified a lever-pressing behavioral task^19^ in which mice choose between either collecting high-value rewards with progressively increasing cost, or low-value rewards with fixed cost. To maximize the reward-to-cost ratio, mice must keep track of the cost changes and update their strategy accordingly. Using this approach, we inferred belief updating in *Grin2a^Y700X+/−^* mice and compared it to wild-type (WT) littermates. We also used computational approaches to quantitatively understand the differences in behavioral patterns and strategies between the two groups during this process.

Converging evidence suggests that the thalamus is altered in schizophrenia patients, including volume, neuron count, metabolism, and connectivity^20–26^. However, mechanistic causal links between thalamic changes and schizophrenia-related symptoms are poorly understood. Using functional ultrasound imaging we found aberrant functional connectivity in the mediodorsal thalamus (MD) in *Grin2a^Y700X+/−^* mice. Combining *in vivo* single unit recording, optogenetic manipulations and behavioral analysis we identified the MD as an important node for keeping track of the environmental changes and allowing mice to form rational behavioral strategies accordingly. More importantly, we demonstrated that reduced neuronal activity in MD is a key pathology underlying belief-driven strategy updating in *Grin2a^Y700X+/−^* mice and enhancing neuronal activity in the MD successfully rescues belief updating deficits in *Grin2a^Y700X+/−^* mice.

### Grin2a^Y700X+/−^ mice exhibited slower belief updating and unstable behavioral states

To validate the knock-in of the *Y700X* mutation of *Grin2a*, we examined the NR2A protein level in the brain of *Grin2a^Y700X+/−^* mice. Western blots showed that knock-in of the *Y700X* mutation significantly decreased the expression of NR2A in the brain of heterozygous mice, while the expression of NR2A was depleted in homozygous mice, validating the knock-in of the mutation (Fig.S1). Since the mutation was found to be heterozygous in human patients, we used heterozygous mice as the mutant experiment group and WT litter-mate mice as the control group. We characterized the phenotypic profile of *Grin2a^Y700X+/−^* mice using a battery of behavioral assays measuring symptom domains often tested in other mouse models for schizophrenia^27,28^, including the open field test (OFT), elevated O-maze test, sucrose preference test (SPT), and pre-pulse inhibition of the startle reflex test (PPI). *Grin2a^Y700X+/−^* mice did not show differences from WT controls in travel distance (Fig.S2a) and time (Fig.S2b) in the OFT, suggesting normal locomotion. They also traveled equally in the center of the open field (Fig. S2c-d) and in the open arm of the O-maze (Fig.S2e and f) compared to WT controls, suggesting no anxiety-like behaviors. We used SPT to measure anhedonia-like phenotype and found no differences in their preference for sucrose solution compared to WT mice (Fig.S2g). We used the pre-pulse inhibition (PPI) of startle response to evaluate sensorimotor gating ^29,30^ and found that *Grin2a^Y700X+/−^*mice didn’t show deficits in either the startle response (Fig.S2h) or the PPI ratio (Fig.S2i).

Next, we trained WT and *Grin2a^Y700X+/−^* mice on a lever-pressing task we adapted^19^ to capture the belief updating process (Fig.1). In this task, a mouse is provided with two levers to choose between on single trials, one that yields high reward (HR lever, 3 drops of milk) and another that yields low reward (LR lever, 1 drop of milk) (Fig.1a). At the beginning of each behavioral session, the HR lever requires one press to deliver reward while the LR requires six. As such, both WT and *Grin2a^Y700X+/−^* mice showed preference to the HR lever: after sampling both sides for a few trials, they would choose HR lever in consecutive trials (Fig.1d and Fig.S3a). Following 4 consecutive HR lever choices, its press requirement was progressively increased (one press every time for each HR trial; Fig.1b). LR lever cost was held constant (Fig.1b). With this design, the optimal strategy to maximize the reward would be choosing HR when the reward/press of the HR lever was higher, while choosing LR after the equal value point (HR required press number=18 for 3 drops of milk) (Fig.1c). Typically, WT mice would show behavior consistent with an optimal strategy, meaning that they showed a high probability of choosing HR lever (p(HR)) when the HR lever press requirement was low, but gradually shifted to the LR lever as the cost increased (Fig.1d-f). We define the mice finishing the shift from HR to LR lever (choosing LR more than 6 times in a row) as one block (Fig.1d-f). On average, p(HR) drops to 0 when the HR required press number equals 22, exceeding the equal value point (Fig.1e, f, and S3b). Most of the blocks (>99%) were finished before the HR required press number reaches 50. *Grin2a^Y700X+/−^* mice were able to make adaptive decisions as the environment changed (Fig.1e and f), but only fully shifted to LR choice in 48% (74 out of 153) of the blocks, and this shift occurred later than in WT mice (Fig.S3b). In 52% of the blocks, mutant mice p(HR) didn’t drop to 0 before the HR required press number reached 50. Indeed, mutant mice had significantly longer blocks than WT (Fig.1g) and reduced optimality (Fig. 1h). These results demonstrated that mutant mice had a slower shift from HR to LR lever when the required press number of HR progressively increased. To examine whether this shift was a result of altered reward or cost sensitivity, we performed two additional experiments. In one case we kept the press number requirements of both levers equal and static but the reward ratio of the two levers at 1:3 (Fig.S4a). In this experiment, it took equivalent trials for WT and mutant mice to commit to the high reward lever (Fig.S4a). In the other case, we kept the rewards equal for both levers while the requirement was higher for one lever (12 vs. 6, Fig.S4b) and found that both groups of mice committed to the low-cost lever after an equivalent number of trials (Fig.S4b). These results suggested that mutant mice were equally-sensitive to cost and reward in a static environment and that the slower shift in decision-making was not due to the inability to detect reward or cost differences. Together, these data suggest that mutant mice exhibit a selective deficit in adapting to dynamic environmental changes.

**Fig.1:**
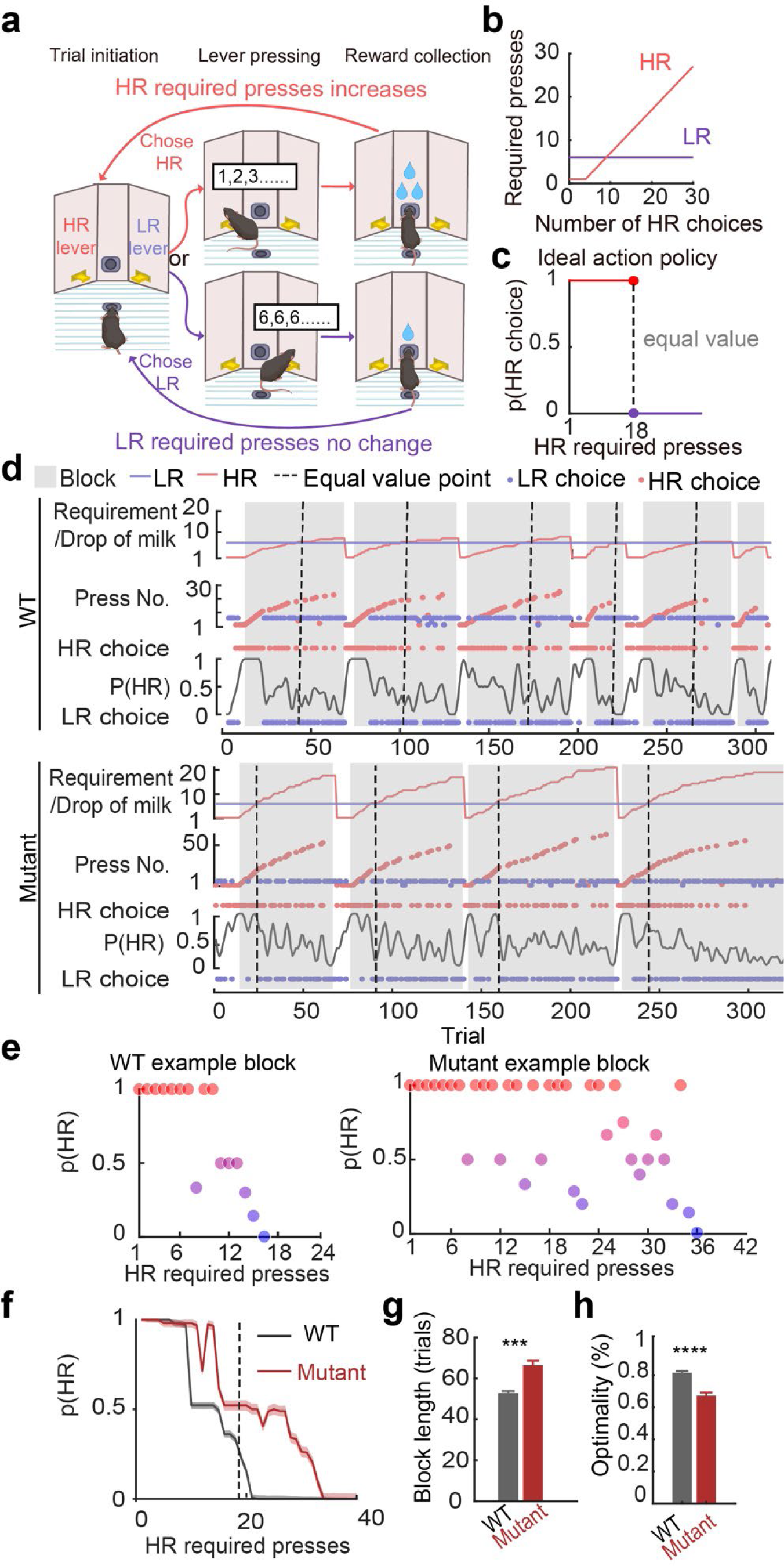
*Grin2a^Y700X+/−^* mice showed suboptimal performance in the belief updating lever-pressing task. **a,** Schematic of the lever-pressing task. Food-restricted mice were required to press levers to collect rewards. On each trial, a mouse freely chooses one of the two levers, leading to high (HR) or low reward (LR). The LR lever provides one drop of milk with a requirement of 6 presses. The HR lever, on the other side, offers three drops of milk, but achieving this high reward requires the pressing numbers that increase with each successful reward. **b,** Pattern of HR and LR cost change. **c,** Ideal action policy to maximize reward per press. **d,** An example session of a WT (top) and mutant (bottom) mouse. First row: Required press number/anticipated drops of milk of HR and LR in each trial; Second row: mouse’s press number of each trial (Red dot: HR choice; Blue dot: LR choice); Third row: mouse’s choice in each trial (Red dot: HR choice; Blue dot: LR choice) and the calculated probability of choose HR (black line). **e,** Probability of choosing HR at each HR requirement of an example block of a WT (left) and mutant (right) mouse. **f,** Summarized plot (median +/− SEM) of WT (n=308 from 9 mice) and mutant (n=153 from 8 mice). **g,** Block length is longer for mutant than WT mice. Kolmogorov-Smirnov test. **h,** Optimality of the performance of mutant mice is significantly lower than WT mice. Kolmogorov-Smirnov test. Optimality =Animal (Reward/presses)/Ideal (Reward/presses). (***, P<0.001; ****, P<0.0001.)

To ask whether the suboptimal behavior of mutant mice was a result of aberrant dynamic belief updating, we built a computational model based on Bayesian inference (Fig.S5) and asked which factors in belief updating were disrupted in *Grin2a^Y700X+/−^* mice. In each trial, the model integrates prior belief with the most recent sensory evidence (a noise-corrupted estimate of the number of presses required on most recent trial) using a fixed belief update rate to generate a posterior belief that gives rise to a decision (Fig.S5a). We fit the model to mouse behavior by adjusting parameters to capture key factors in the belief updating process, belief updating rate parameter (α), the initial uncertainty on HR beliefs (*σ*_*HR*_), the fixed uncertainty associated with LR beliefs (*σ*_*LR*_) and the degree of noise associated with cost perceptions (*δ*). The belief updating rate parameter controls the relative weighting between prior and current perceived evidence, *σ*_*HR*_ and *σ*_*LR*_control the width (and thus uncertainty) of belief distributions, and the perceptual noise parameter (*δ*) controls the scale of noise in cost perception. Malfunctions in any factor can disrupt belief updating process. The fitted models for WT and *Grin2a^Y700X+/−^* can closely mirror the behavior observed in WT mice and *Grin2a^Y700X+/−^* mice respectively (Fig.S5b). The parameters of the fitted model that capture the *Grin2a^Y700X+/−^* mice performance have a lower belief update rate (lower α, indicating weighing more on the prior) (Fig.S5c) while showing no difference in other parameters. The model suggested that the slower shift from HR choice to LR choice in the *Grin2a^Y700X+/−^* mice is not caused by noisy perception but is mainly a result of the slow belief updating rate (Fig.S5d).

We noticed that mice demonstrated different behavioral strategies adapting to the environmental changes during the lever-pressing task. We observed three distinct behavioral states: one when mice tended to consistently choose HR (HR-committed state), one when mice consistently chose LR (LR-committed state), and another one when mice constantly switched between HR and LR sides (exploration state). To further understand these behavioral patterns, we applied an unsupervised Hidden Markov Model (HMM) to quantitively detect the proportions and transition probabilities of the three behavioral states in each block (Fig.2a). With HMM, we were able to assess the likelihood of each state for every trial. In this context, the state assigned to each trial corresponds to the one with the highest probability (Fig.2b). We found that both WT and *Grin2a^Y700X+/−^* mice started with HR-committed state, switched to the exploration state, and then the probability of LR-committed state rose as the value of HR decreased (Fig.2b). For WT mice, in each trial, there was a greater likelihood of remaining in the current state rather than transitioning to the next one in the subsequent trial (Fig.2c). This suggested that each state was stably maintained. *Grin2a^Y700X+/−^* mice were able to maintain the stability of the HR-committed state similarly to WT mice, but when they entered exploration state, they had a higher probability of switching to LR-commitment (4% for WT v.s. 18% for mutant). More strikingly, even when they entered the LR-committed state, there was a near 100% probability they would go back to the exploration state, thus switching back and forth between the exploration and LR-commitment state (Fig.2c). This also led to longer exploration of mutant mice compared to WT (Fig.2d). These results showed that mutant mice had unstable behavioral states and therefore exhibited longer exploration states of each block (Fig.2c and d).

**Fig.2:**
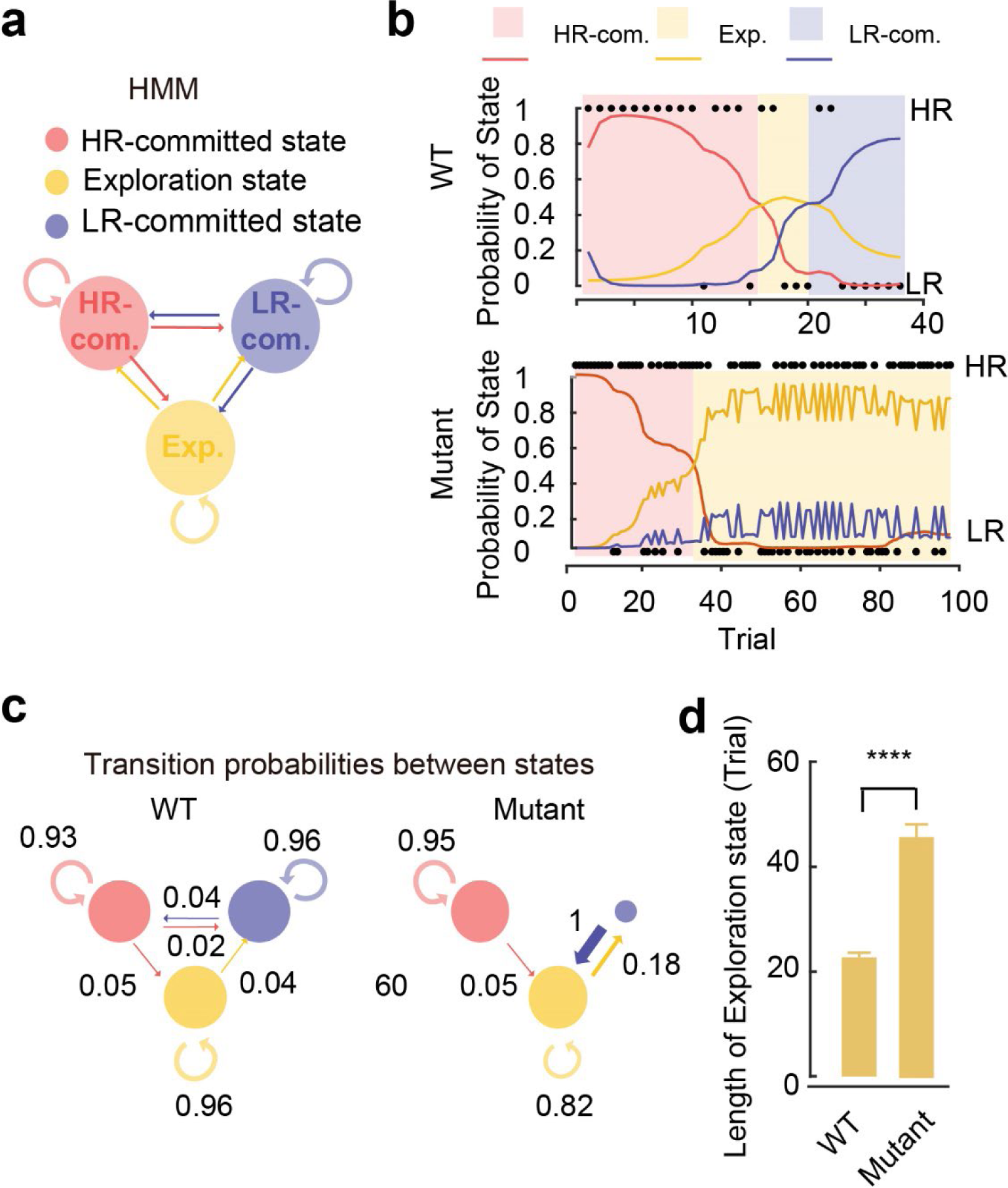
HMM model reveals Unstable behavioral states of *Grin2a^Y700X+/−^* mice. **a**, Schematic of the HMM. **b,** probability of each state at each trial of one example block from WT (top) and *Grin2a^Y700X+/−^* mice (bottom). The shaded area indicated the identified behavioral states by the HMM (the state with the highest probability in each trial) **c,** Transition probabilities were shown by the area of the circle and the thickness of the arrows. **d,** Trial numbers of the exploration state of WT and *Grin2a^Y700X+/−^* mice. Kolmogorov-Smirnov test. (****, p<0.0001).

### *Grin2a^Y700X+/−^* mice exhibit aberrant activity in MD

To understand the neural substrates underlying the behavioral deficits of *Grin2a^Y700X+/−^* mice, we performed hemodynamic functional ultrasound imaging (fUS) scanning ^31^ on both the WT (n=9) and *Grin2a^Y700X+/−^* mice (n=10) to look for potential regions that show different activities between the groups. We opened a chronic cranial scanning window (anterior to posterior: −2.5mm to 3.00mm; Lateral: −1.5mm to 1.5mm) for each animal. To minimize the movement artifact, we anesthetized the mice and scanned the cranial window for each mouse along the sagittal axis (Fig.3a). We first did pixel-by-pixel local functional connectivity analysis of the whole scanned area. We clustered the pixels based on the differences between WT and *Grin2a^Y700X+/−^* group (characterized by Cohen’s d) and geographical location and identified local clusters with larger Cohen’s d values and larger connected areas. Our data showed larger-area clusters of pixels that showed larger differences (P<0.01), characterized by Cohen’s d, between WT and the *Grin2a^Y700X+/−^*group mainly lies in the dorsal anterior part of the prefrontal cortex (PFC), mediodorsal thalamus (MD) and ventral striatum (vStr) (Fig.3b). To further validate this result, we used the three regions (prelimbic cortex (PL), MD, and vStr) as regions of interest (ROI) and performed seed-based analysis (Fig.3c-h, Fig.S6). Our data revealed that MD had lower dynamic functional connectivity (see methods; Fig.3c and f) in mutant mice. In addition, there was a trend of lower MD-PL functional connectivity in mutants using MD as seed (Fig. S6a, P<0.05 before p-value correction for multiple comparisons and P<0.1 after correction). Similarly, PL to MD functional connectivity tend to be lower in mutant mice (Fig. S6c, P<0.05 before p-value correction for multiple comparisons and P<0.1 after correction). MD seed analysis did not reveal any MD-vStr functional connectivity differences between mutant and WT mice (Fig. S6a). These connectivity changes are strikingly similar to those found in schizophrenia patients ^21^. On the contrary, seeding the vStr or PL did not reveal significant changes from WT littermates (Fig. 3d-h, Fig. S6b and c). These results showed reduced local MD functional connectivity, which covaries with brain activity and indicates information processing efficiency in the area ^32^. This aberrant local functional connectivity was not found in other regions in anesthetized animals. This is consistent with studies on schizophrenia patients showing decreased volume, cell numbers, and metabolism in the MD^22–24,26^.

**Fig.3:**
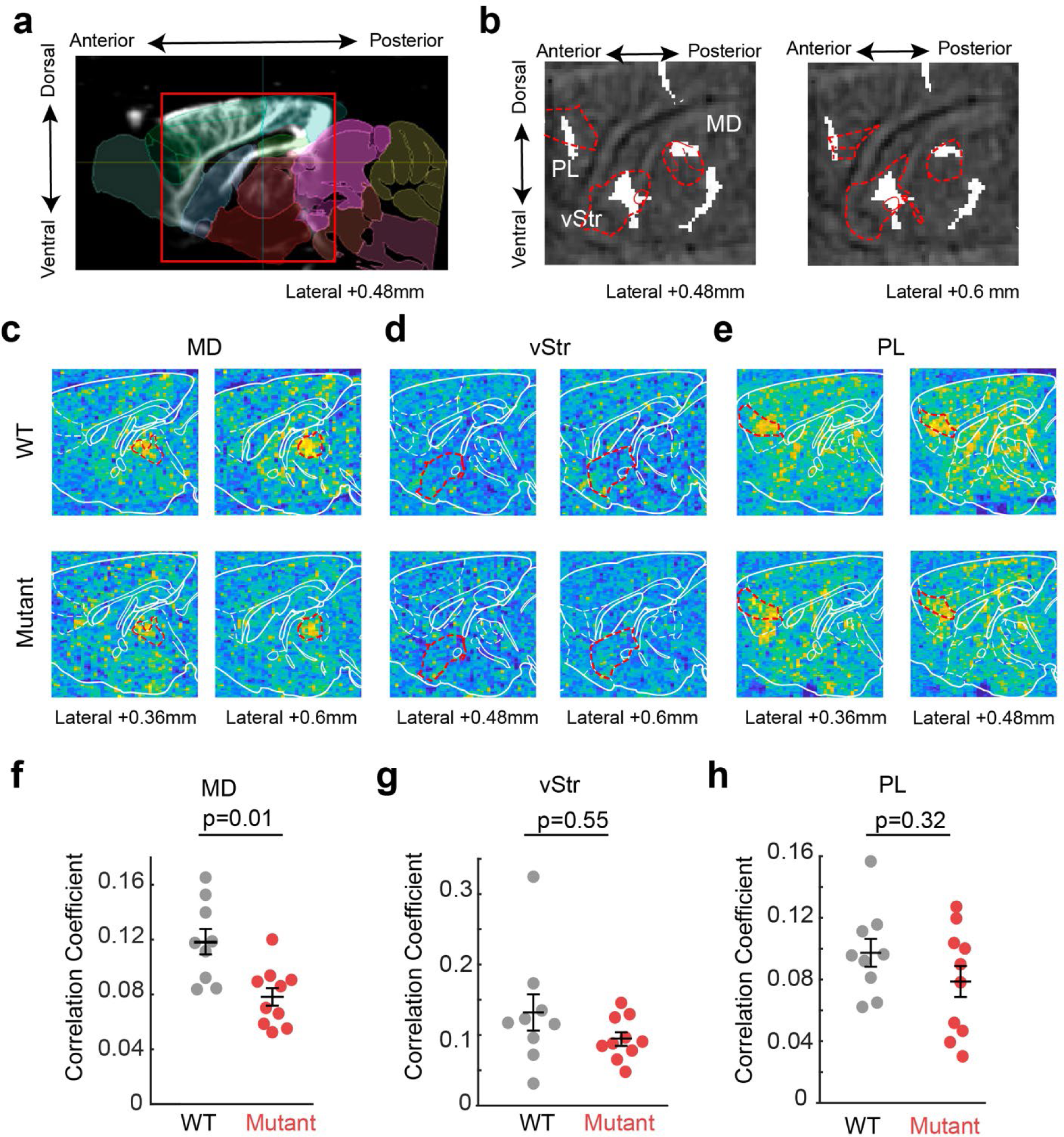
functional ultrasound (fUS) imaging reveals aberrant activity in MD of *Grin2a^Y700X+/−^* mice. **a,** Example of the raw data of one brain slice aligned to the atlas during the fUS scanning. The red rectangle is the scanned area. **b,** pixel-to-pixel comparison of WT and *Grin2a^Y700X+/−^* mice revealed a few regions potentially have distinct activity between the two groups (clusters where Cohen’s d >0.6 and pixel number > 425. Random shuffled test shows the likelihood of forming such clusters is less than 0.01). PL, ventral striatum (vStr), and MD are outlined in red. **c,** Correlation coefficient of two example sagittal brain slices from one WT and one *Grin2a^Y700X+/−^* mice (top and bottom panel correspondingly) taking MD as the seed (seeds are outlined by redline in the heat map). ventral Striatum, and PL as the seed. **d,** Correlation coefficient of two example sagittal brain slices from one WT and one *Grin2a^Y700X+/−^* mice (top and bottom panel correspondingly) taking ventral Striatum (vStr) as the seed. **e,** Correlation coefficient of two example sagittal brain slices from one WT and one *Grin2a^Y700X+/−^*mice (top and bottom panel correspondingly) taking the prelimbic cortex (PL) as the seed. **f,** MD-MD connectivity is significantly lower in *Grin2a^Y700X+/−^* mice compared to WT mice. Unpaired t-test with Welch correction. **g,** vStr-vStr connectivity is not significantly different in *Grin2a^Y700X+/−^* mice compared to WT mice. Unpaired t-test with Welch correction. **h,** PL-PL connectivity is not significantly different in *Grin2a^Y700X+/−^* mice compared to WT mice. Unpaired t-test with Welch correction.

### MD neurons encode dynamic values and behavioral states in the lever-pressing task

As the resting-state fUS imaging revealed that the most significant activity changes in *Grin2a^Y700X+/−^* mice was in MD, and *Grin2a^Y700X+/−^* mice showed compromised performance in the lever-pressing task, we examined whether MD neuronal activity plays an important role in the task. MD has been reported to allow commitment to the new best options when the environment changes in monkeys^33,34^ and the thalamic local field potential correlates with values in dynamic environment in humans^35^. However, how MD neurons operate at the single neuron level in a dynamic environment is an open question. Therefore, we implanted multi-tetrode drives to record neuronal firings from MD while animals were engaged in the lever-pressing task (Fig.S7a). We analyzed 109 MD neurons recorded from 4 WT mice (See methods). We aligned the neuronal activity of each trial to the initiation and reward collection and focused on the signals after reward collection and before the initiation, during which the mice were not actively engaged in pressing levers so that we could study the encoding of the value in the absence of actively collecting the information. We used linear regression to determine whether neuronal responses during any 2s window before initiation encode the HR required press numbers (HR requirement), upcoming choices, and past choices (Fig.4 and Fig.S7). One example MD neuron that fulfilled the criteria of encoding the HR requirement is shown in Fig.4a-c. In total, 82.6% of the neurons encoded the dynamic HR requirement (Fig.4d, Fig.S7b), while far fewer neurons predicted the upcoming choice (44%) or encoded the past choice (41%) (Fig.S7a and b). The percentage of neurons that represent HR requirement and the correlation coefficient of the regression analysis steadily rose at reward collection (Fig.S7b) and reached the peak around 2s before the initiation (Fig.4d and Fig.S7b and S7c), suggesting the most task-related activity in MD appears to be at 2s before the initiation in the task. Next, we used the MD population activity to decode the HR requirement and found that MD population activity was also able to discriminate the high and low HR requirement trials with >70% accuracy (Fig.4e). These results suggest that MD neurons represent the gradually changing effort requirement in the dynamic environment, which is consistent with results from human studies ^35^.

**Fig.4:**
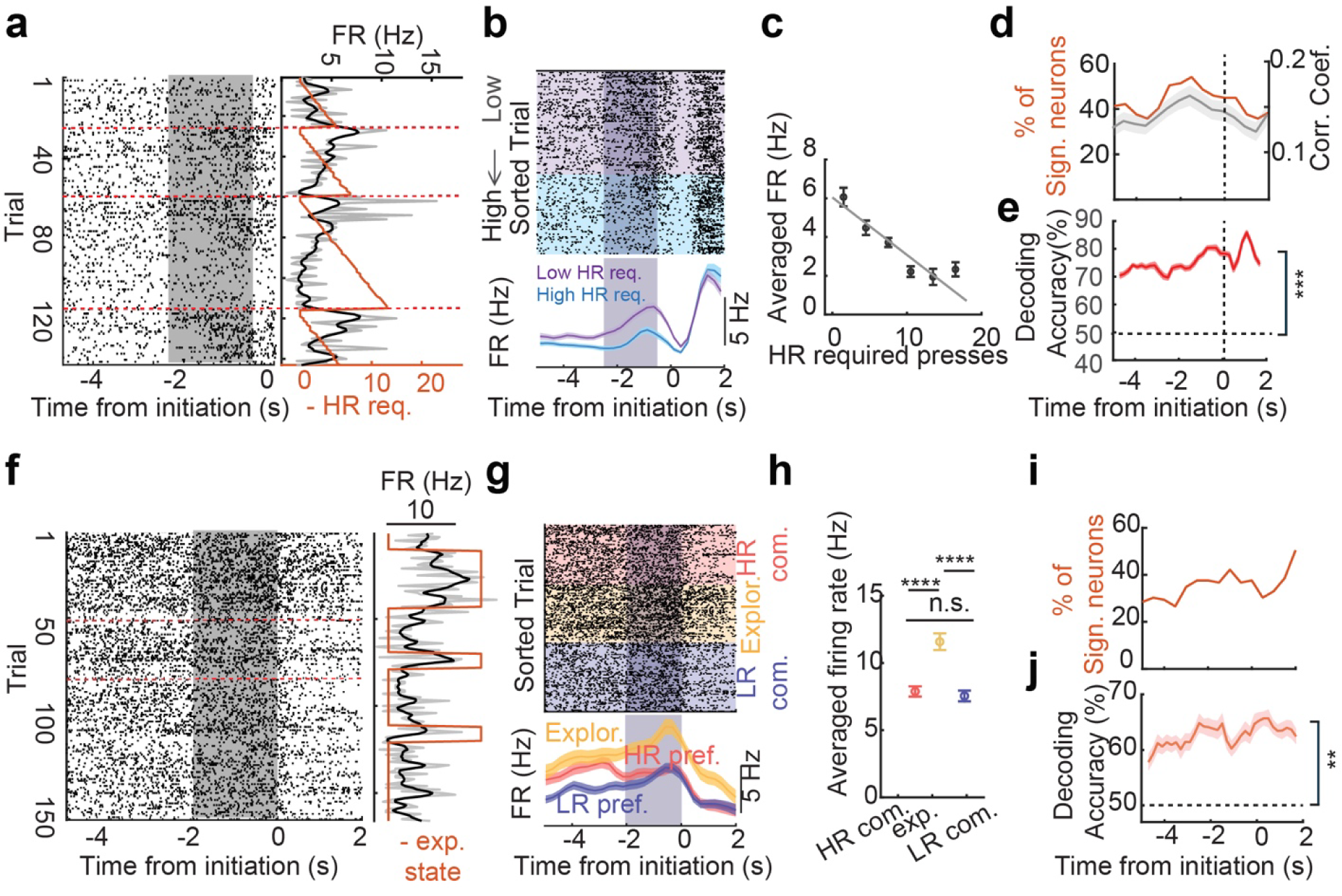
MD neurons encode dynamic values and behavioral states in the lever-pressing task. **a-c,** Example MD neuron whose task activity significantly correlates with upcoming HR cost. a, Left: Raster plot of the example neuron aligned to the initiation, sorted by trial; Right: averaged firing rate (gray), smoothed firing rate of the shaded time window, and the HR required presses(orange) in the corresponding trial on the left. b, Top: Raster plot of the same neuron, sorted by HR required press numbers of the corresponding trial; Bottom: averaged firing rate of low HR requirement trials and high HR requirement trials. FR, firing rate. Req., required presses. c, Regression analysis of the averaged FR with the HR required press numbers of the corresponding trial. R-square=0.264, p=3.21e-11; **d,** Percentage of neurons of which FR significantly correlates with HR required press numbers from MD and correlation coefficient of MD neurons at each time point aligned to initiation (grey line: mean; grey shade: +/−SEM). **e,** Decoding accuracy of the MD population activity in discriminating low/high HR requirement trials. Horizontal dashed line indicates the random decoding accuracy. Z-test. **f-h,** An example neuron from MD showed a significantly different firing rate at the exploration behavioral state compared to HR or LR committed states. f, left: Raster plot of the example neuron plotted aligned to initiation and sorted by trial. Right: averaged firing rate (gray), smoothed firing rate (black) of the shaded time window, and the exploration state (orange) in the corresponding trial on the left. g, Top: Raster plot of the same neuron from f sorted by behavioral states. Bottom: Averaged FR of the same neuron at corresponding behavioral states. h, Averaged FR of the same neuron at different behavioral states. One-way ANOVA test with Tukey’s multiple comparisons. **i,** Percentage of neurons of which FR during the exploration is significantly different from the committed state in MD at different time points. **j,** Decoding accuracy (Data are mean+/−SEM) of all recorded MD neurons for exploration v.s. Committed states. Z-test. (****, p<0.0001, n.s., p>0.05; **, p<0.01; ***, p<0.001)

Since *Grin2a^Y700X+/−^*mice also showed unstable behavioral states, we then asked whether MD neuron activity helps stabilize behavioral states. To answer this question, we applied the HMM model to the behavioral data from our electrophysiological recordings. With the HMM model, we categorized trials into HR- or LR-committed state and exploration state trials (Fig.4F-j and Fig.S7d). We found that 75.2% of MD neurons have distinguished firing rates in different states before the initiation. Fig. 4f-h represents one neuron that discriminates the exploration state and the committed state. The encoding of the behavioral states of MD neurons occurred during the period throughout the reward collection to the initiation (Fig. 4i and Fig. S7d). Further, we used the MD population activity to predict the exploration state from the committed state, and we found that MD activity can decode the states with higher accuracy (more than 60%) than random (50%, Fig. 4j and Fig.S7e).

### Inhibition of MD in WT mice phenocopies Grin2a^Y700X+/−^ mice

Since MD neurons encode HR requirements and behavioral states during the lever-pressing task, the impairment of MD neuronal activity will potentially lead to dysfunctional evaluation and suboptimal choices when the environment changes, as well as disrupted behavioral states. To test how loss of function manipulation of MD activity affects the belief updating, we injected AAV2/9-CamkII-iC++-mCherry bilaterally in MD and implanted optical fibers above the injection site (Fig. 5a). When mice were performing during the lever-pressing task, we shed blue light to inhibit neuronal activity after the mice finished one trial and terminated the light illumination once the initiation was completed. We randomly alternated the light-on and off blocks in each session. Our results showed that inhibition of MD in WT mice during the lever-pressing task slowed down the decrease of p(HR) when HR requirements were progressively increased (Fig. 5b), which led to the increased block length (Fig. 5c). This slowed adaptation to the changing environment also led to decreased optimality (Fig. 5d). Next, we applied HMM to the behavioral data collected from the optogenetically manipulated mice. We found that with MD inhibition, mice could still form a stabilized HR-committed state when the block started, indicated by a high probability of staying at the HR-committed state (Fig. 5f). However, when the animals entered the exploration state, they had a higher probability of transitioning to the LR-committed state (50% v.s. 4%) in the laser-on blocks compared to the laser-off blocks (Fig. 5f). When the mice entered the LR-committed state, they also had a much higher chance of switching back to the exploration state in the laser-on blocks (90% v.s. 0) (Fig.5 e-f). This switching back and forth led to unstable behavioral states and a longer exploration state (Fig. 5g). This result indicates that MD neuronal activity is critically important for mice to keep track of the changing required effort and form a stable behavioral state in a dynamic environment. This result also showed that the WT mice with MD inhibition phenocopied the performance of the *Grin2a^Y700X+/−^* mice: a high chance of switching back and forth between the LR-committed state to the exploration state and longer exploration state.

**Fig.5:**
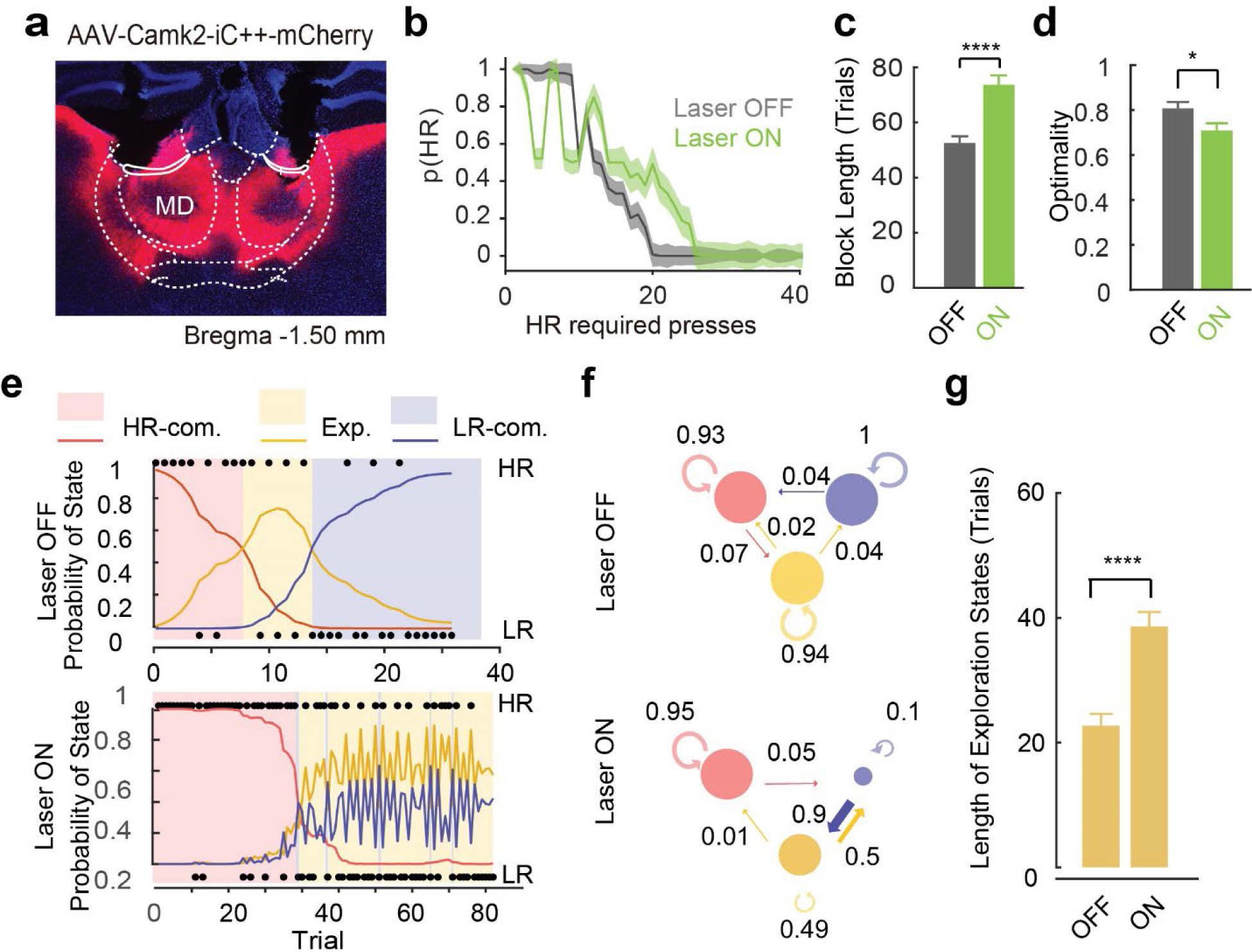
Inhibition of MD neural activity in WT mice phenocopies the performance of *Grin2a^Y700X+/−^* mice in the lever-pressing task. **a,** Virus injection site of optogenetic inhibition test. Red: AAV-Camk2-iC++-mCherry; Blue: Dapi. **b,** Summarized performance of MD inhibition (n=72 for light OFF and n=66 for light ON from 7 mice). data are median +/− SEM. **c,** Block length of laser OFF and ON sessions. unpaired t-test. **d,** Optimality of light off and on sessions. Kolmogorov-Smirnov test. **e,** HMM detection of the probability of behavioral states at each trial of an example block of MD light OFF (top) and ON (bottom). **f,** The HMM model detected the probability of the transition between states of all MD OFF blocks (top) and ON blocks (bottom). **g,** The length of exploration state of all blocks without and with MD inhibition. Kolmogorov-Smirnov test. (****, p<0.0001; *, P<0,05; n.s. p>0.05).

### Activation of MD rescued impaired performance of Grin2a^Y700X+/−^ mice in the lever-pressing task

With the observation that inhibition of MD showed a similar phenotype of *Grin2a^Y700X+/−^*mice: sub-optimal behavioral choices, longer block length, and switching back and forth from exploration state and LR-committed state, we hypothesized that the aberrant MD activity of *Grin2a^Y700X+/−^*mice was a cause of the impaired performance in the lever-pressing task, and we would be able to rescue the performance of *Grin2a^Y700X+/−^*mice in the lever-pressing task by enhancing neuronal activity in MD. To test this hypothesis, we injected AAV2/9-Camk2-ChR2(C12S/D156A)-mCherry in bilateral MD in mutant mice (Fig.6a) and trained these mice on the lever-pressing task. ChR2(C12S/D156A) is a stabilized step function opsin (SSFO), and the cation channel opens with a brief blue light illumination and ends with orange light illuminations^36^. This channel enables subthreshold depolarization of neurons for a prolonged period and this method can be used to enhance the MD function^36,37^. After 3 weeks of virus expression, we optically activated MD at the beginning of a session and terminated the activation at the end of a session in the lever-pressing task. We randomly alternated laser on and off sessions so that we could compare the behavior with and without MD activation of the same mutant mice (53 blocks for OFF and 45 blocks for ON). Our data showed that with MD activation, the shift from HR levers to LR levers in *Grin2a^Y700X+/−^* mice became faster (Fig.6b). Along with the earlier shift, the block length was significantly decreased (Fig.6c) and the optimality of the behavioral performance was increased (Fig.6d). Thus, MD activation led to a significant enhancement of behavioral performance in mutant mice. We then applied the HMM model to the MD-activated and control conditions to investigate the behavioral states of each group. We found that *Grin2a^Y700X+/−^*mice with virus expression in MD but laser off showed similar behavioral state patterns compared to *Grin2a^Y700X+/−^* mice without virus injection (Fig. 2b): switching back and forth between exploration state (Fig.6e-f) and longer exploration; However, when we turned on the light to activate MD, the *Grin2a^Y700X+/−^* mice were capable to maintain stabilized states (Fig.6e-f) and had shorter exploration states (Fig.6g). These results showed that MD activation in the *Grin2a^Y700X+/−^*mice could rescue the behavioral deficits in the lever-pressing task.

**Fig.6:**
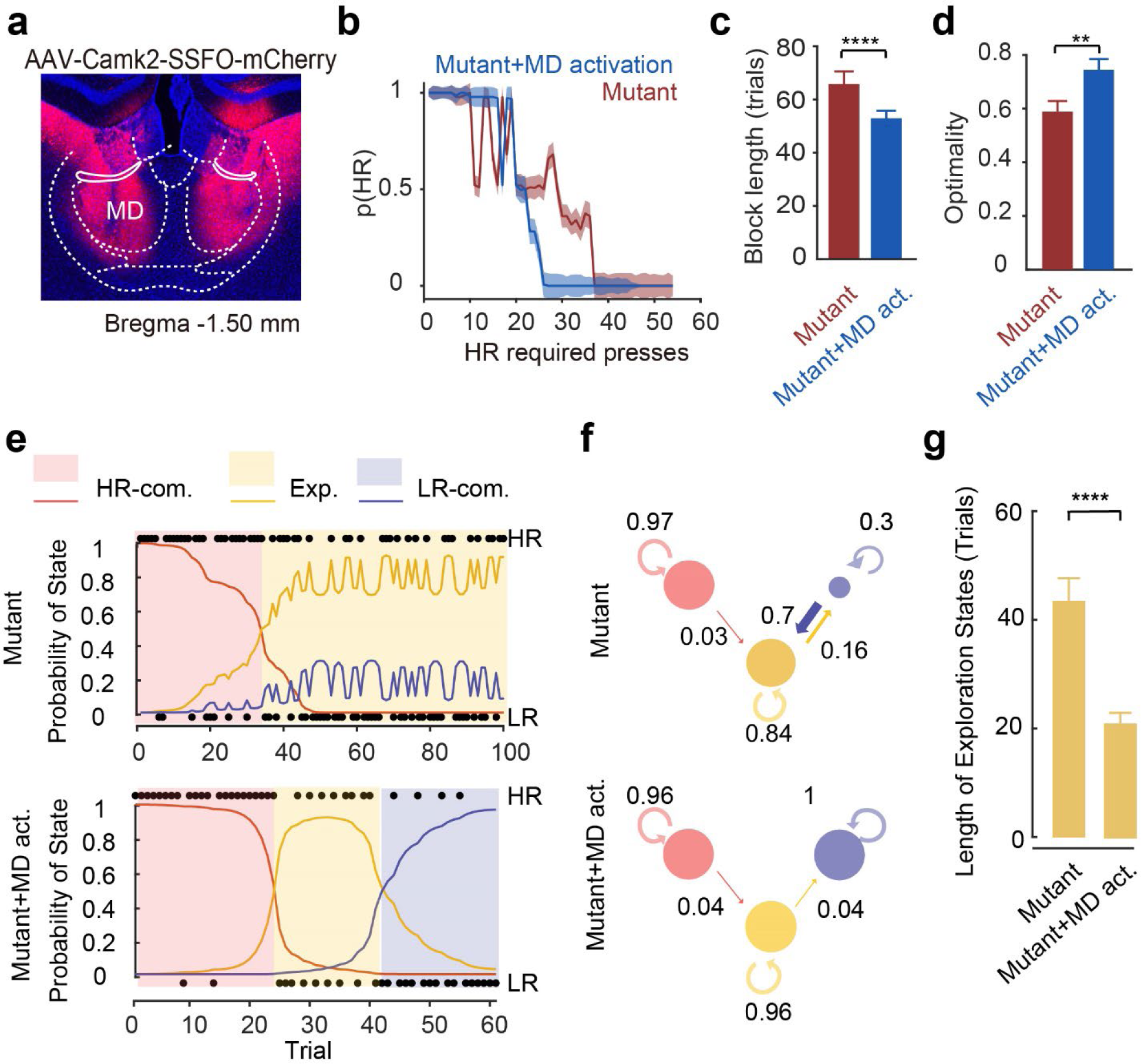
Activation of MD neural activity restores the performance in *Grin2a^Y700X+/−^* mice in the lever-pressing task. **a,** Virus injection site of SSFO activation of MD during the lever-pressing task. Red: AAV-Camk2-chR2(C128S/D156A)-mCherry; Blue: DAPI. **b,** MD activation with SSFO enhanced the behavioral performance of mutant mice (n=53 blocks of mutant, and n=45 blocks of mutant +MD activation group from n=5 mice); Data are Median +/− SEM; **c,** block length is shorted with MD activation. unpaired t-test; **d,** Optimality is improved with MD activation. Kolmogorov-Smirnov test. **e,** Example blocks of a mutant mouse without MD activation (top) and with MD activation (bottom) **f,** MD activation with SSFO reversed the unstable behavioral states in mutant mice. (Top: sessions without MD activation; Bottom: sessions with MD activation.) **g,** MD activation decreased the length of exploration in the *Grin2a^Y700X+/−^*mice. Kolmogorov-Smirnov test. (****, p<0.0001; **, P<0.01; ***., p<0.001).

## Discussion

In this study, we established a mouse model with a patient-derived loss-of-function mutation (Y700X) of the Grin2a gene identified in a large-scale exome sequencing study ^2^. We also developed a quantitative behavioral paradigm for mice, the lever-pressing task with dynamic value changes, to monitor the belief updating process, the impairment of which has been proposed to be the computational mechanism of psychosis in humans ^14,16–18,38,39^. Our behavioral and modeling data indicate that *Grin2a^Y700X+/−^* mice exhibited a slower belief update rate and unstable behavioral states in the lever-pressing task. Functional ultrasound imaging results showed reduced local functional connectivity in MD, consistent with human studies showing impaired MD structure and/or function in schizophrenia patients ^21,23,24,26^. *In vivo* single-unit electrophysiological recording and optogenetic inhibition experiments revealed that MD neuronal activity tracks environmental changes, which is in line with earlier studies showing that MD neural activity tracks context ^40^. Elevating the MD neuronal activity in *Grin2a^Y700X+/−^* mice rescued the impaired belief updating and unstable behavioral states in the lever-pressing task. Taken together, our work reliably models a type of cognitive dysfunction akin to that seen in patients with schizophrenia and provides tools to study underlying neural mechanisms. Our finding that MD plays an important role in the cognitive deficits in the schizophrenia model could also open the door for exploring novel therapeutic targets and strategies.

It is well established that schizophrenia has a substantial genetic component. Though genome-wide association studies (GWAS) have identified 287 common risk loci of schizophrenia^41^, these genetic variations have small effects individually and are predominantly non-coding. One of the most robustly associated genetic factors is the copy number variations (CNVs) such as the 22q11.2 deletion. Mouse models carrying 22q11.2 deletions have been widely studied and has significant advanced our understanding of the neurobiological effects of this deletion and its relevance to schizophrenia^42–44^. Although the CNVs generate large effects, they disrupt multiple genes simultaneously, which will limit our ability to derive clear functional insights^28^. With the collective efforts from the SCHEMA consortium, loss-of-function mutations from 10 genes including Grin2a were identified as ultra-rare variations that confer substantial risk for schizophrenia^2^. These findings provide an unprecedented opportunity to model and study molecular, cellular and circuit changes relevant to schizophrenia pathology. The finding of *Grin2a* loss of function mutations is consistent with the long existing glutamate hypofunction hypothesis of schizophrenia pathology. Indeed, mice with *Grin2a* genetic mutations have been used as a model for schizophrenia based on this hypothesis long before the current genetic evidence. While in humans, the variations of the *Grin2a* gene were found in the form of point mutations in heterozygous patients, most of the behavioral phenotypes for loss of function in Grin2a were found in NR2A full knockout mice^45,46^. The robust cognitive phenotype in heterozygous *Grin2a^Y700X+/−^* mice provides an opportunity to dissect neurobiological mechanisms in a system more closely modeling the genetic risks of schizophrenia.

Computational psychiatry approaches provide mathematical models that can explain the core symptoms of schizophrenia^18,38,47,48^, the effort taking advantage of these models has seen major progress in modeling schizophrenia-related symptoms in rodents^49^. For example, in the work of modeling hallucination-like perception in mice, it was shown that hallucination-like perception in mice can be explained by stronger prior in Bayesian belief updating model^49^. Taking this route, we developed a quantitative behavioral paradigm as a readout of the belief updating process. In this behavioral paradigm, we created a progressively changing environment, through which we can monitor how mice make adaptive decisions. The mainstream behavioral paradigms for studying belief updating in mice usually use abrupt environmental changes, such as reward value change indicated by a cue^50^, change of probability of reward delivery in block structure^51^ or reversal of rules^52,53^. Fundamentally, our task also takes advantage of environmental change to study belief updating; however, instead of an abrupt change of option values in the task, the cost of one option in the task was gradually changing while the other one remained constant. This structure allows us to monitor not only how fast the animals update their beliefs, but also the strategy and behavioral state changes during the belief updating process, which is a direct readout of the animals’ estimation of the uncertainty of the environment. With this design, we were also able to detect how early the mice started to sample the other option, suggesting the mice’s internal evaluation of the uncertainty level of the environment. Our finding that *Grin2a^Y700X+/−^* mice started to sample early in the progressively changing environment (Fig. S3c) indicated that these mice overestimated uncertainty compared to WT. These results are consistent with findings in human research showing that higher uncertainty levels were perceived in people with psychosis^54–58^.

Our functional ultrasound imaging results revealed a localized neural activity impairment in MD, and we showed that neuronal activity is critical for optimal belief updating process. Consistent with this idea, we found that enhancing MD activity could rescue belief updating in *Grin2a^Y700X+/−^* mice. A growing body of work have shown that MD plays a major role in cognitive functions such as working memory^37,59^ task switching^40,60^ and decision making under uncertainty^61–63^, which are known to be affected in schizophrenia^6^. Interestingly, the expression of *Grin2a* is not limited to MD, and it remains to be discovered how mutations of *Grin2a* severely affected MD activity. Transcriptomic and proteomic studies showed molecular evidence for hypofunction in the prefrontal cortex in the heterozygous *Grin2a* knock out mice^64^. Since MD is bidirectionally connected with the prefrontal cortex and *Grin2a* is highly expressed in this area, one possibility could be that lack of NR2A impaired the functional inputs from the prefrontal cortex to MD and thus affecting the functional maturation of MD during development ^37,53,64,65^.

Schizophrenia is a mental disorder with multiple risk factors and multiple affected functional domains. Here we only studied one genetic risk factor and one aspect of the symptoms. It is still not known whether different risk factors will converge on the same pathway to mediate common symptoms. With the identification of loss-of-function mutations in 10 high risk genes by the large-scale SCHEMA study^2^, it will be important to systematically investigate in a set of animal models potential common phenotypes and converging mechanisms at molecular, cellular, and circuit levels^43,66^. Such studies may bring us one step closer to understanding schizophrenia pathophysiology and underlying neurobiological mechanisms.

## Author Contribution

R.D.W. and R.L. developed the lever-pressing task in coordination with M.M.H. T.Z. performed most of the behavior, functional ultrasound imaging, in vivo electrophysiological recording, optogenetic manipulation experiments with the help of YY.H. and K.H. and mentorship from M.M.H. and G.F.. T.Z. performed behavioral data analysis with mentorship from M.M.H. T.Z. wrote the manuscript with feedback from YY.H., K.H., N.H., A.F. M.N., and M.M.H.. YY.H. did the data analysis for functional ultrasound imaging and in vivo electrophysiological recordings with help from J.S. and mentorship from M.M.H. and built HMM and belief updating models with mentorship from M.N. and M.M.H. K.H. performed the reward and cost sensitivity test and analyzed the data with help from E.H.. A.F. and K.H. performed OFT, O-maze, and SPT experiments and analyzed the data. C.L. performed the western blot experiment and analyzed the data. J.W. and X.G. generated the *Grin2a ^Y700X+/−^* mouse model. Tarjinder Singh shared genetic variants information in schizophrenia patients.

## Acknowledgement

The work was supported by NIMH grant R01MH134466 to MMH and GF. This work in Feng lab was also supported by funding from the Poitras Center for Psychiatric Disorders Research at MIT, the K. Lisa Yang and Hock E. Tan Center for Molecular Therapeutics in Neuroscience of the Yang Tan Collective at MIT, the Stelling Family Research Fund at MIT, and the Stanley Center for Psychiatric Research at Broad Institute of MIT and Harvard. T.Z. receives funding support from the Brain and Behavior Research Foundation (BBRF #31216). We thank the SCHEMA consortium for sharing genetic variants information in schizophrenia patients. We thank Guillermo Horga for his insightful comments on the manuscript. We thank Frederico Azevedo and William Menges for their help with functional ultrasound data analysis.

---

**Fig. S1:**
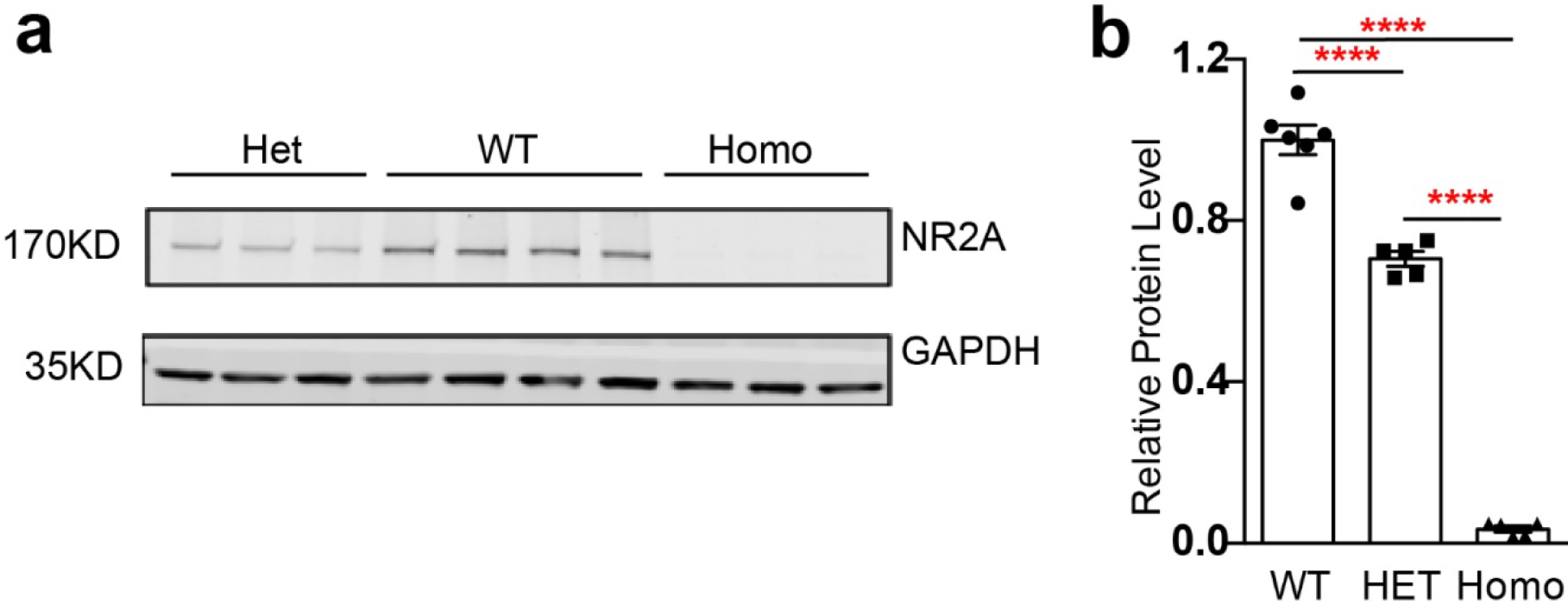
Western blot showed the decrease of NR2A levels of *Grin2a^Y700X+/−^* mice. **a,** Example of one run of western blot experiments of the whole brain tissue of WT, Het (*Grin2a^Y700X+/−^*), and Homo (*Grin2a^Y700X+/+^*) mice. **b,** Summary of the relative NR2A protein level in WT, Het, and homozygous mice. (****, P<0.0001, one-way ANOVA with Tukey’s multiple comparisons test data from 6 mice from WT, 5 mice from Heterozygous and 5 mice from homozygous).

**Fig. S2.**
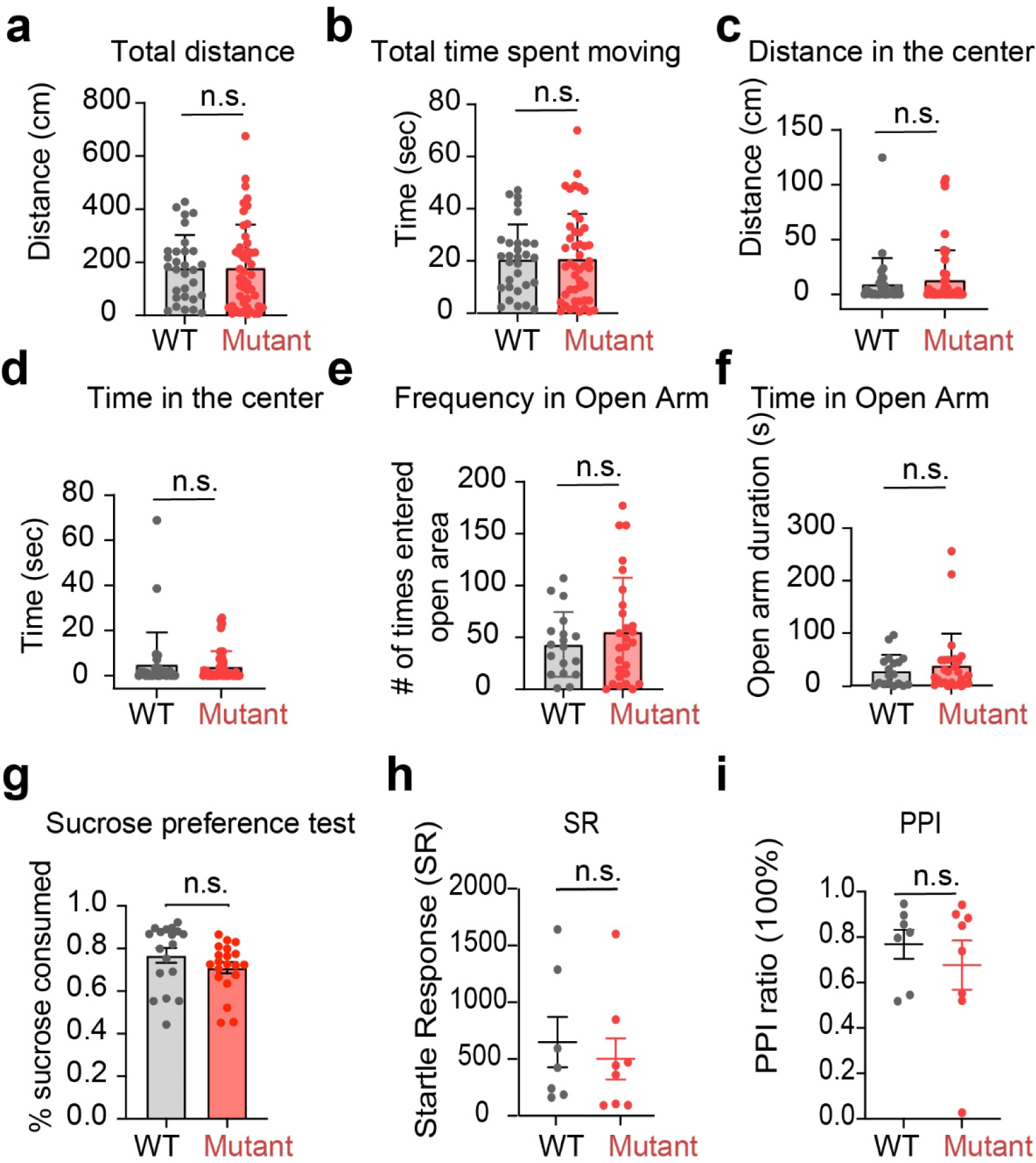
*Grin2a^Y700X+/−^* mice didn’t show deficits in locomotion, anxiety, sucrose preference, and pre-pulse inhibition test. **a** and **b**, Total distance and total time spent moving in the open field test of WT and mutant mice are not different. n=30 for WT and n=47 mutant. Unpaired t-test. **c** and **d**, Distance and total time spent in the center of the open field of WT and mutant are not different. Kolmogorov-Smirnov D test. **e** and **f**, Frequency and time spent in the open arm in the O-maze test are not different for WT and mutant mice. Kolmogorov-Smirnov D test. n=20 WT and n=30 mutant. Unpaired t-test. **g,** The preference for sucrose in the sucrose preference test is not different for WT and mutant mice. n=18 for WT and n=20 for mutant. Kolmogorov-Smirnov test. **h,** The pre-pulse inhibition ratio is not different for WT and mutant mice. n=7 for WT and n=8 for mutant. Unpaired t-test. (n.s., P>0.05)

**Fig. S3.**
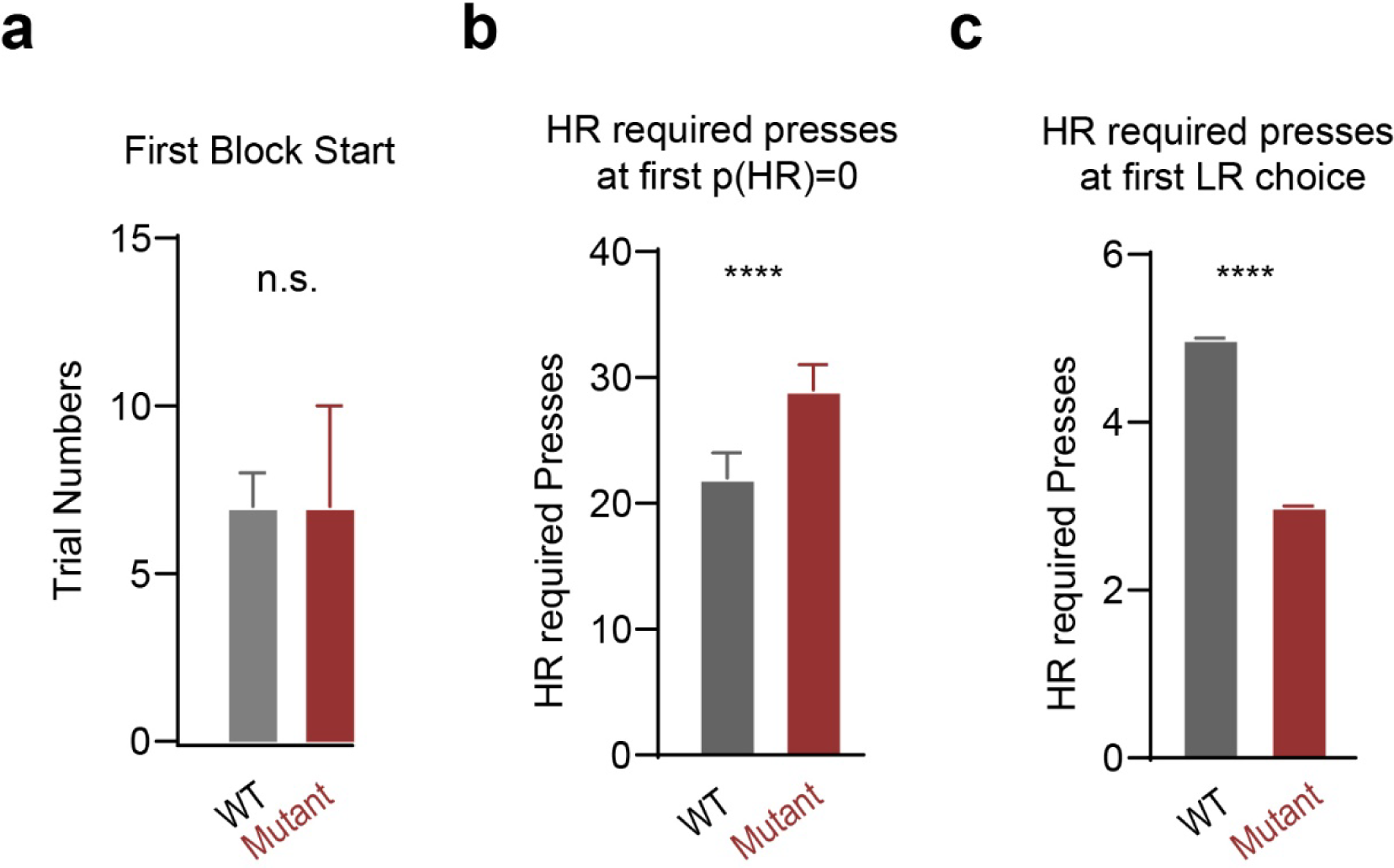
*Grin2a^Y700X+/−^* mice started sampling earlier and stopped sampling later than WT mice. **a,** Trial numbers that it took mice to start consecutively choosing HR lever once the mice started each session. Median with 95% CI. Kolmogorov-Smirnov test (n=160 from 9 mice for WT. n=97 from 8 mice for Mutant). **b,** Distribution of the HR required press numbers at p(HR)=0, i.e., mice fully shift to the LR side, of WT and Mutant mice. Kolmogorov-Smirnov test. (n=308 from 9 mice for WT and n=74 from 8 mice for mutant (only 74 blocks from 153 blocks reached p(HR)=0 before HR required presses reaches 50.)) **c,** Distribution of the HR required press numbers at the first LR choice in a block of WT and Mutant mice. Kolmogorov-Smirnov test (n=308 from 9 mice for WT and n=153 from 8 mice for mutant).

**Fig. S4.**
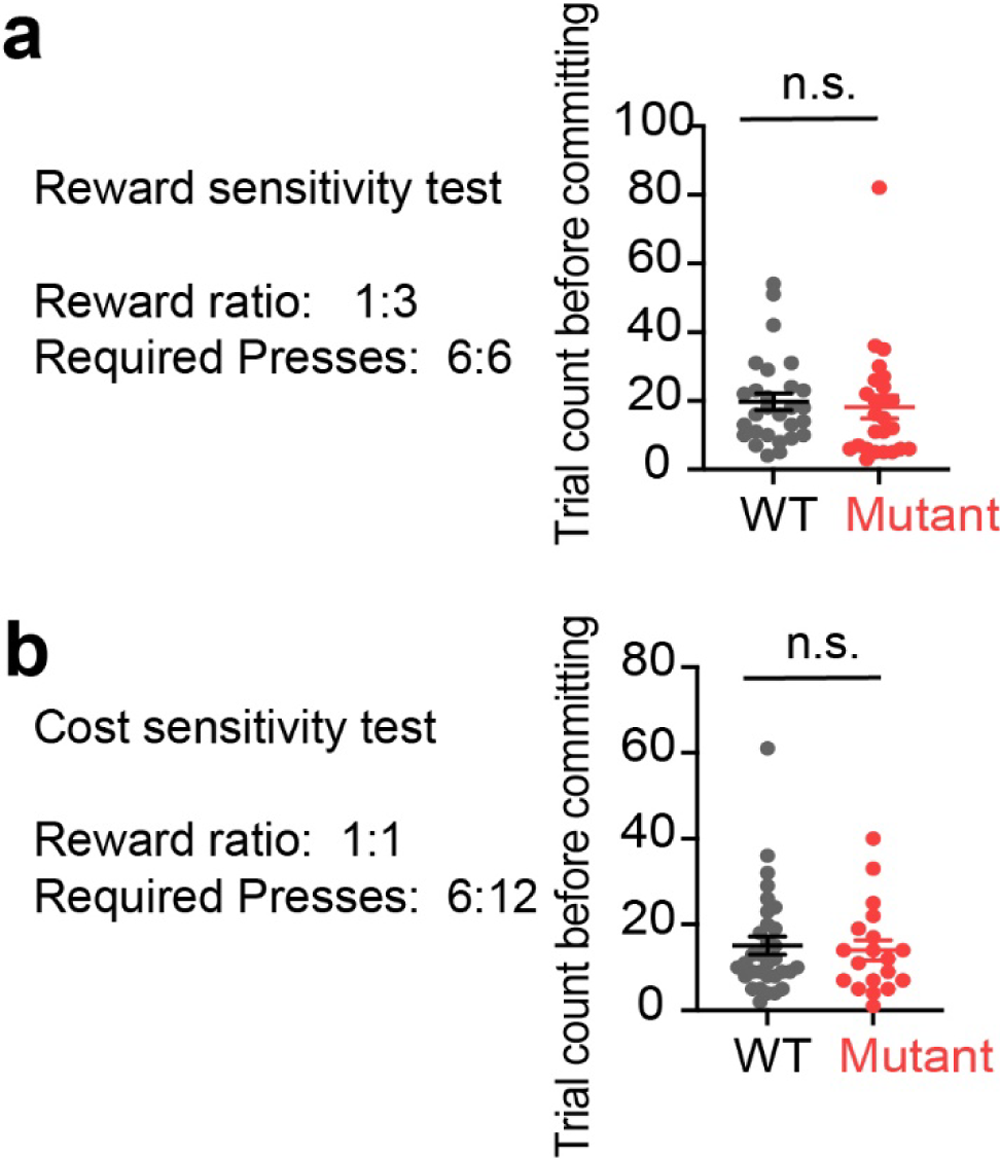
*Grin2a^Y700X+/−^* mice can detect cost and reward differences in a static environment. **a,** Reward ratio was set as 3:1 on the HR side and the LR side while the required press numbers of both sides are 6. The trial numbers it takes the mice to fully commit to the HR side (5 consecutive choosing HR) is not different between the WT and mutant group. Unpaired t-test. **b,** Reward was equal on both sides of the lever, while one side of the lever required the pressing number to be 6 while the other side 12. The trial numbers the mice took to fully commit to the low requirement side are not different between the WT and mutant groups. Unpaired t-test. (n.s. p>0.05; WT: n=28 from 6 mice; Mutant: n=19 from 6 mice)

**Fig. S5.**
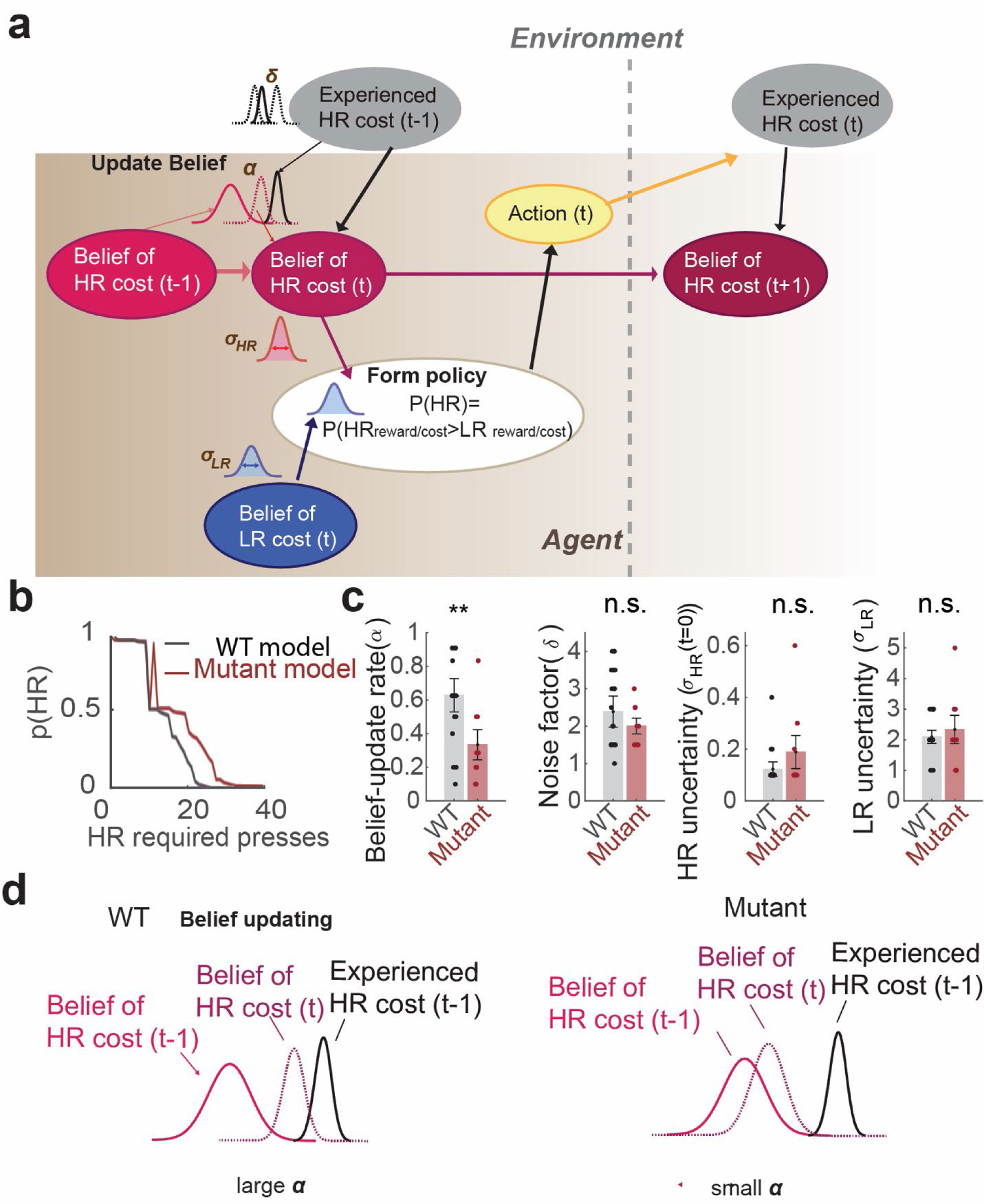
Belief updating model accounts for the impaired behavioral performance of *Grin2a^Y700X+/−^* mice. **a,** Schematic of the belief updating model. The belief of the HR cost of trial t is derived from the Bayesian inference model and policy is formed based on the probability of the relative value of the two choices. **b,** Models fitted with WT and mutant mice performance can qualitatively capture the performance of each group. **c,** Model parameters fitted to the WT and mutant empirical data show that mutant mice have a lower belief update rate. **d,** Schematic demonstration of the belief update process in WT and mutant mice, where mutant agent shows slower belief updating rate and puts higher weights on prior.

**Fig. S6.**
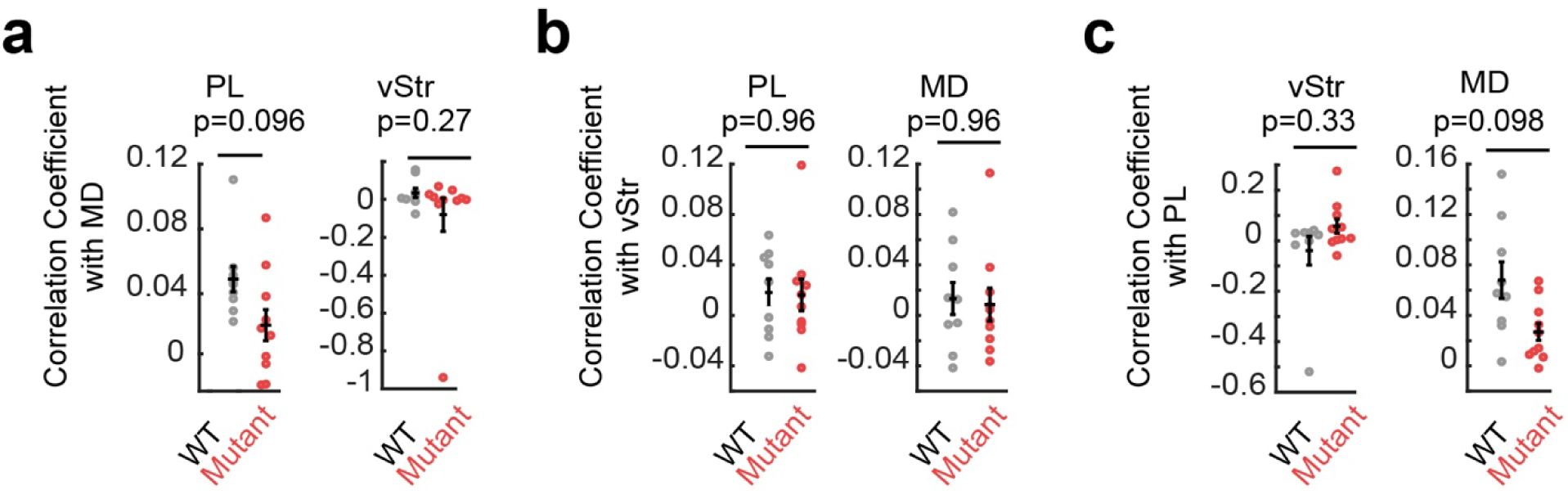
Functional ultrasound imaging showed the whole brain connectivity of *Grin2a^Y700X+/−^* mice. **a,** Correlation coefficient with PL and vStr using MD as seed are not different between WT and *Grin2a^Y700X+/−^* mice. Unpaired t-test with Welch correction. **b,** Correlation coefficient with PL and MD using vStr as seed are not different between WT and *Grin2a^Y700X+/−^* mice. Unpaired t-test with Welch correction. **c,** Correlation coefficient with vStr and MD using PL as seed are not different between WT and *Grin2a^Y700X+/−^* mice. Unpaired t-test with Welch correction. (n.s., p>0.05; *, p<0.05)

**Fig. S7.**
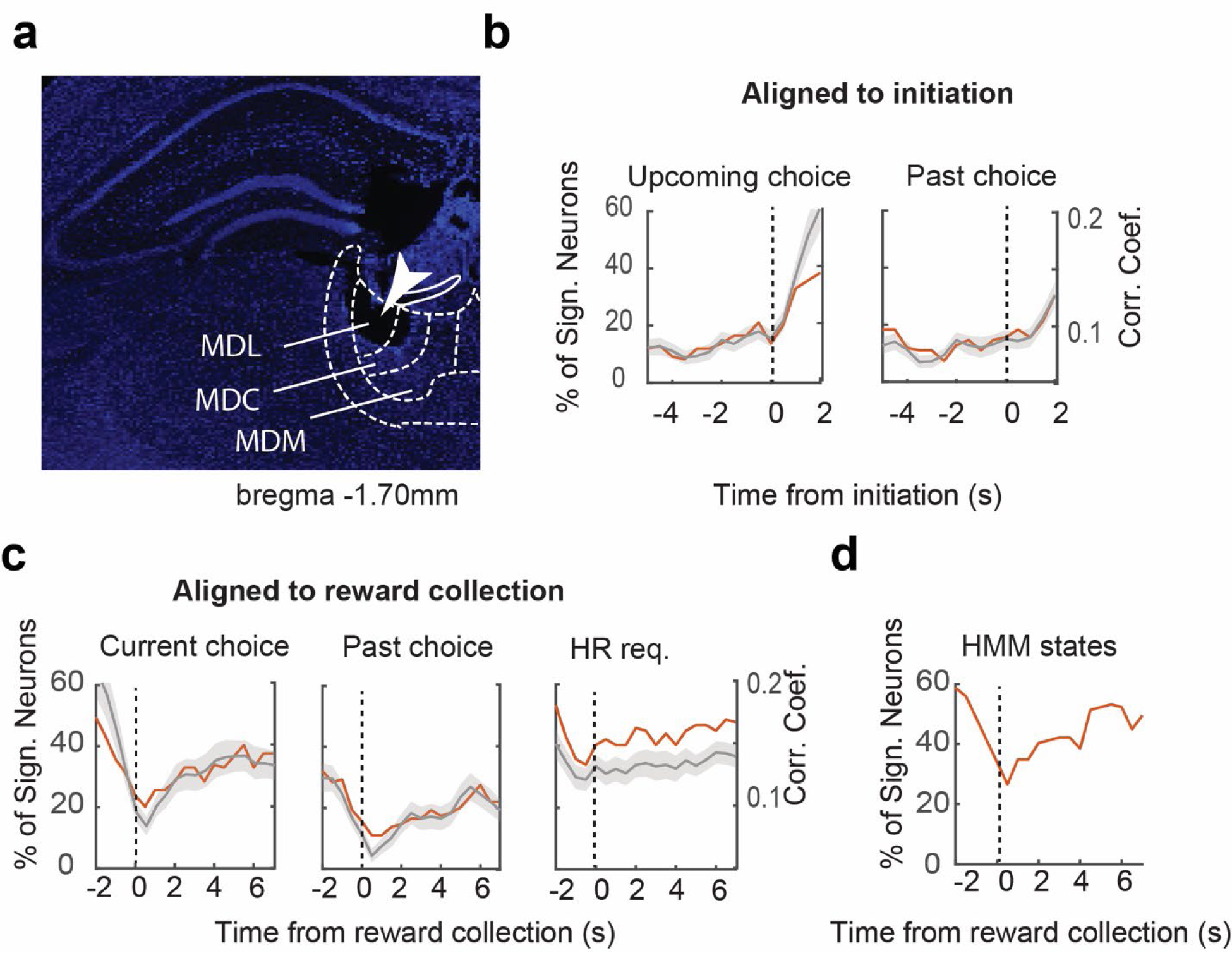
Neural activity in MD encodes HR required presses from reward collection to the initiation of the next trial, but shows higher choice encoding before reward collection and after initiation. **a,** Histology showing example lesion in the MD resulting from passing current through tetrodes before euthanasia. The white arrow indicates the lesion by current injection and the tetrode trace. MDL, lateral MD. MDC, central MD. MDM, medial MD. **b,** Percentage of neurons of which firing rate (FR) significantly correlates with the choice of the upcoming trial (top) or the choice of the past trial (bottom) from MD and the correlation coefficient of all neurons from MD at each time point aligned to initiation (grey line: mean; grey shade: mean+/−SEM). **c,** Percentage of neurons of which FR significantly correlates with the choice of the upcoming trial (left), the choice of the past trial (middle), or the HR required presses (right) from MD and the correlation coefficient of all neurons from MD at each time point aligned to Reward collection (grey line: mean; grey shade: mean +/−SEM). Note that the encoding of the current choice increases after the reward collection, it could reflect the reward volume difference associated with the choice. **d,** Percentage of neurons of which the FR is significantly different at exploration state and committed state at each time point aligned to reward collection. Req, required press numbers.

