## Supplementary figures and methods for "Enhancement of mediodorsal thalamus rescues aberrant belief dynamics in a mouse model with schizophrenia-associated mutation"

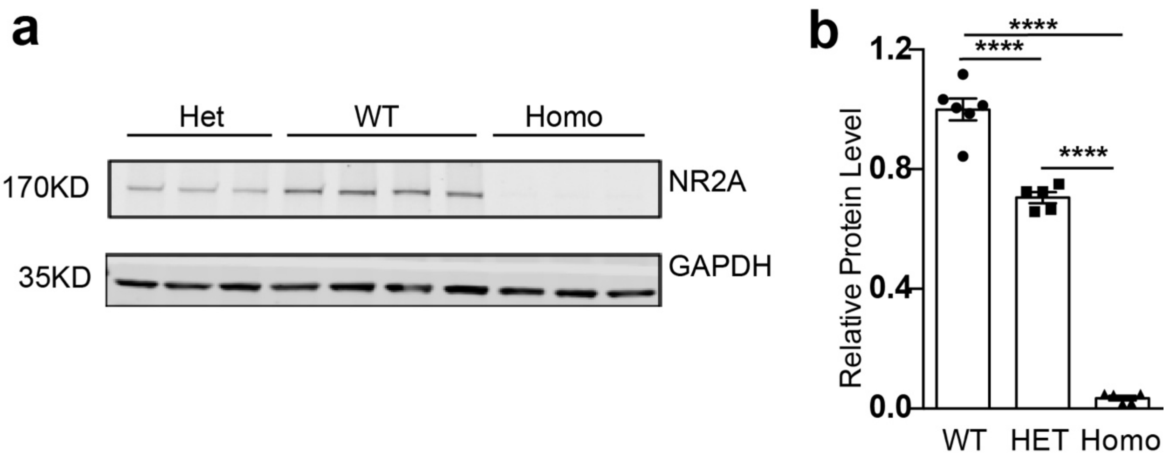


**Fig. S1: Western blot showed the decrease of NR2A levels of *Grin2a*$^{Y700X+/-}$ mice.**
**a,** Example of one run of western blot experiments of the whole brain tissue of WT, Het (*Grin2a*$^{Y700X+/-}$), and Homo (*Grin2a*$^{Y700X+/+}$) mice. **b,** Summary of the relative NR2A protein level in WT, Het, and homozygous mice. (****, $P<0.0001$, one-way ANOVA with Tukey's multiple comparisons test data from 6 mice from WT, 5 mice from Heterozygous and 5 mice from homozygous).

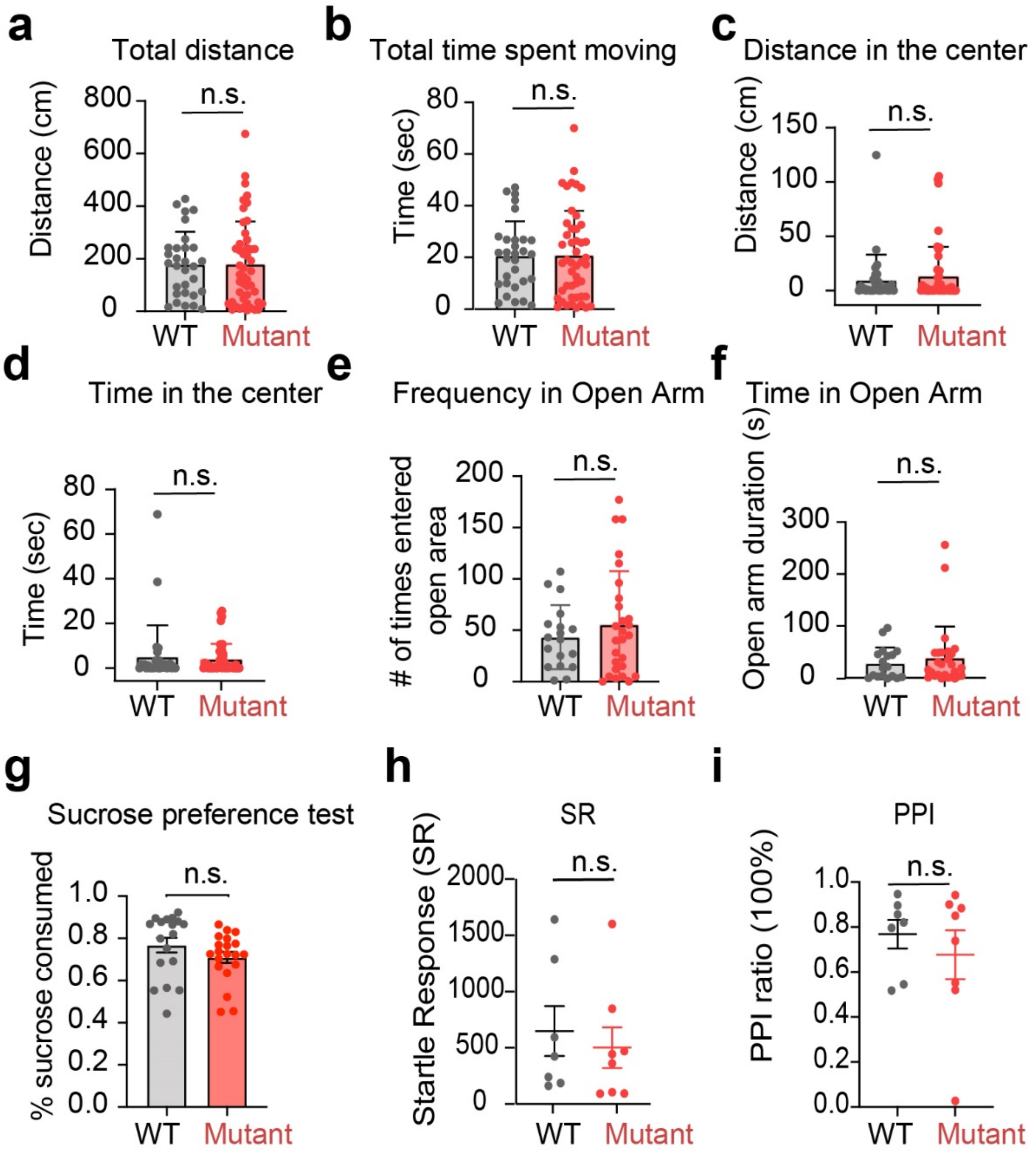


**Fig. S2 *Grin2a*$^{Y700X+/-}$ mice didn't show deficits in locomotion, anxiety, sucrose preference, and pre-pulse inhibition test.**

**a** and **b**, Total distance and total time spent moving in the open field test of WT and mutant mice are not different. n=30 for WT and n=47 mutant. Unpaired t-test. **c** and **d**, Distance and total time spent in the center of the open field of WT and mutant are not different. Kolmogorov-Smirnov D test. **e** and **f**, Frequency and time spent in the open arm in the O-maze test are not different for WT and mutant mice. Kolmogorov-Smirnov D test. n=20 WT and n=30 mutant. Unpaired t-test. **g,** The preference for sucrose in the sucrose preference test is not different for WT and mutant mice. n=18 for WT and n=20 for mutant. Kolmogorov-Smirnov test. **h,** The pre-pulse inhibition ratio is not different for WT and mutant mice. n=7 for WT and n=8 for mutant. Unpaired t-test. (n.s., P>0.05)

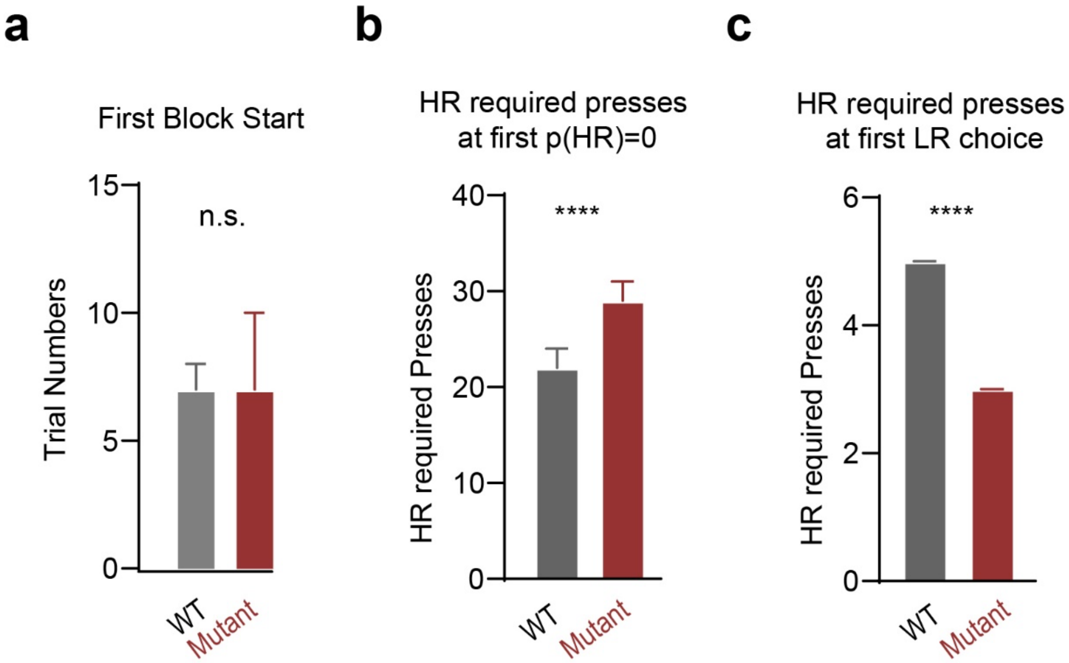


**Fig. S3 *Grin2a*$^{Y700X+/-}$ mice started sampling earlier and stopped sampling later than WT mice.**

**a,** Trial numbers that it took mice to start consecutively choosing HR lever once the mice started each session. Median with 95% CI. Kolmogorov-Smirnov test (n=160 from 9 mice for WT. n=97 from 8 mice for Mutant). **b,** Distribution of the HR required press numbers at p(HR)=0, i.e., mice fully shift to the LR side, of WT and Mutant mice. Kolmogorov-Smirnov test. (n=308 from 9 mice for WT and n=74 from 8 mice for mutant (only 74 blocks from 153 blocks reached p(HR)=0 before HR required presses reaches 50.)) **c,** Distribution of the HR required press numbers at the first LR choice in a block of WT and Mutant mice. Kolmogorov-Smirnov test (n=308 from 9 mice for WT and n=153 from 8 mice for mutant).

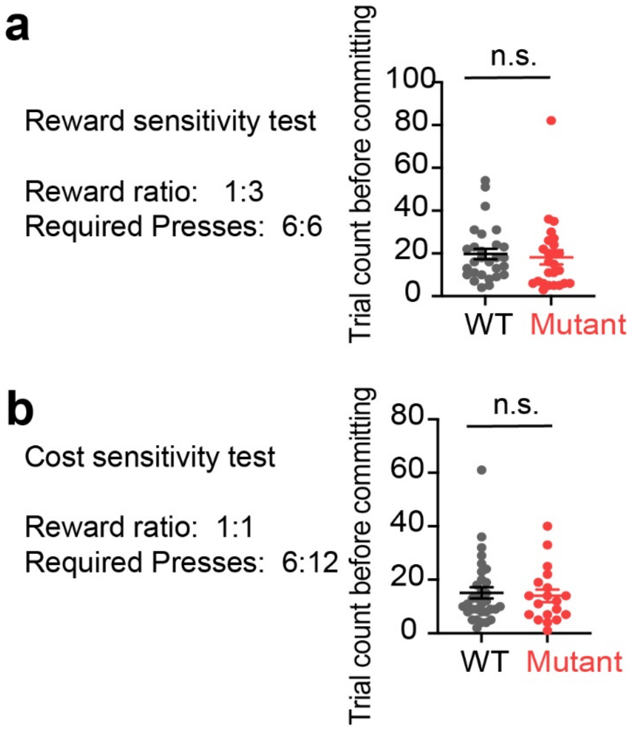


**Fig. S4 *Grin2a*$^{Y700X+/-}$ mice can detect cost and reward differences in a static environment.**
**a,** Reward ratio was set as 3:1 on the HR side and the LR side while the required press numbers of both sides are 6. The trial numbers it takes the mice to fully commit to the HR side (5 consecutive choosing HR) is not different between the WT and mutant group. Unpaired t-test. **b,** Reward was equal on both sides of the lever, while one side of the lever required the pressing number to be 6 while the other side 12. The trial numbers the mice took to fully commit to the low requirement side are not different between the WT and mutant groups. Unpaired t-test. (n.s. $p>0.05$; WT: n=28 from 6 mice; Mutant: n=19 from 6 mice)

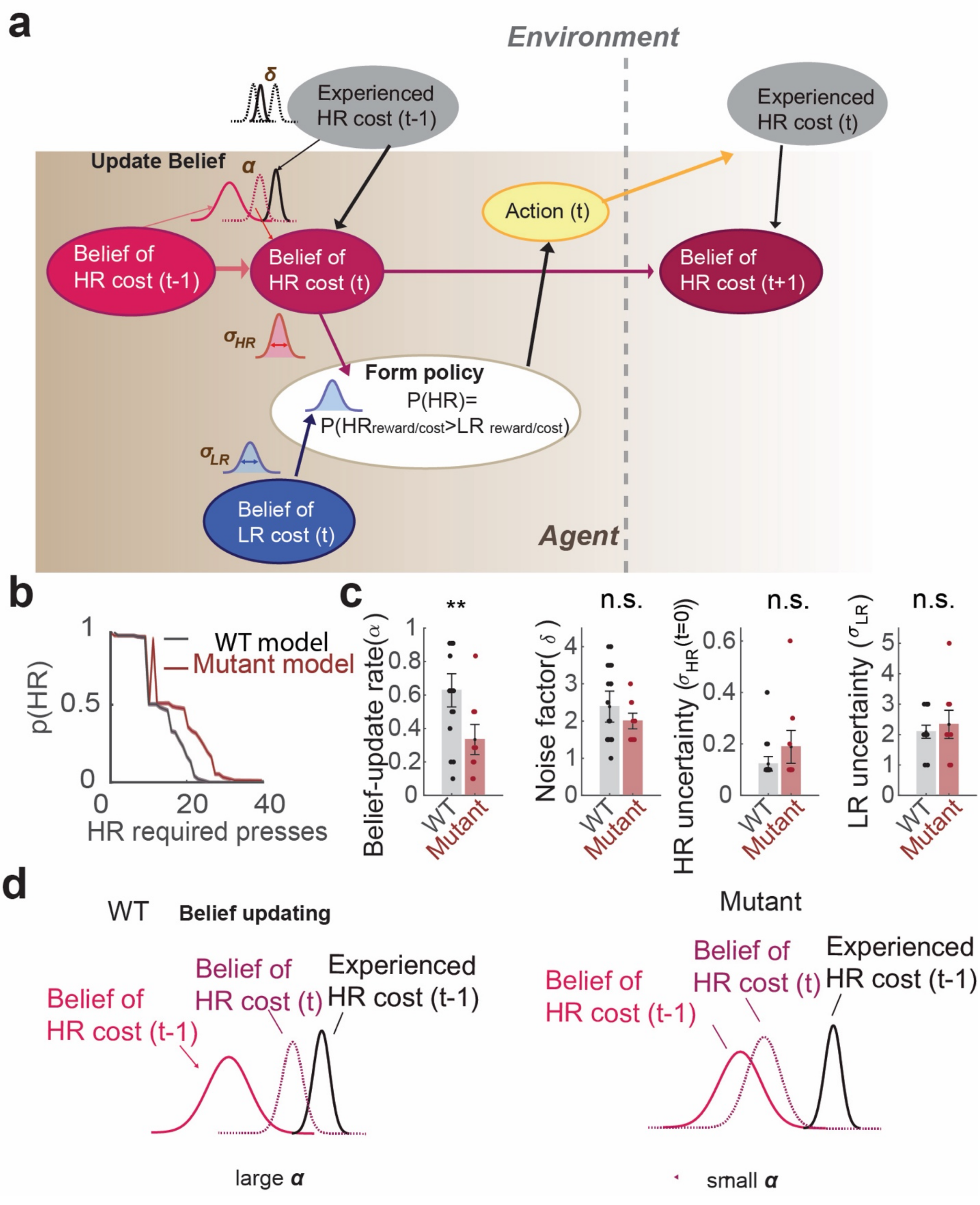


**Fig. S5 Belief updating model accounts for the impaired behavioral performance of *Grin2a*$^{Y700X+/-}$ mice.**

**a,** Schematic of the belief updating model. The belief of the HR cost of trial t is derived from the Bayesian inference model and policy is formed based on the probability of the relative value of the two choices. **b,** Models fitted with WT and mutant mice performance can qualitatively capture the performance of each group. **c,** Model parameters fitted to the WT and mutant empirical data show that mutant mice have a lower belief update rate. **d,** Schematic demonstration of the belief update process in WT and mutant mice, where mutant agent shows slower belief updating rate and puts higher weights on prior.

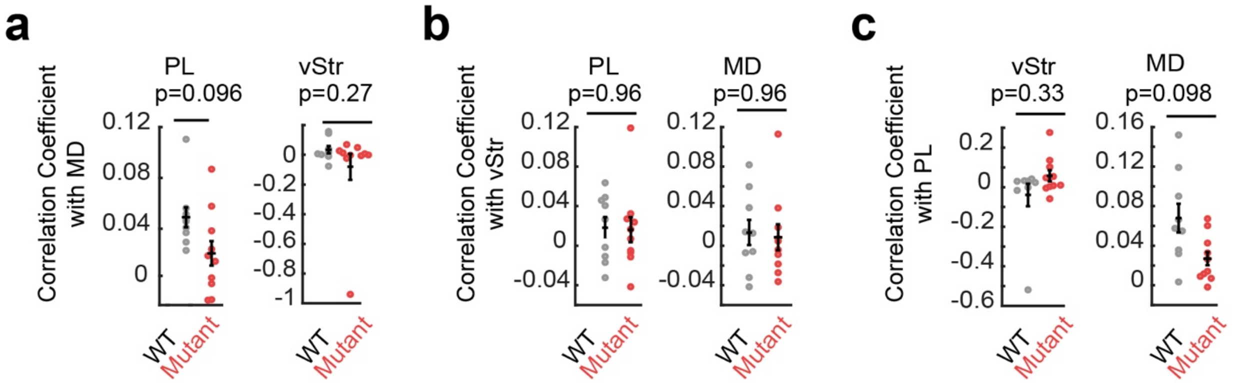


**Fig. S6 Functional ultrasound imaging showed the whole brain connectivity of *Grin2a*$^{Y700X+/-}$ mice.**

**a,** Correlation coefficient with PL and vStr using MD as seed are not different between WT and *Grin2a*$^{Y700X+/-}$ mice. Unpaired t-test with Welch correction. **b,** Correlation coefficient with PL and MD using vStr as seed are not different between WT and *Grin2a*$^{Y700X+/-}$ mice. Unpaired t-test with Welch correction. **c,** Correlation coefficient with vStr and MD using PL as seed are not different between WT and *Grin2a*$^{Y700X+/-}$ mice. Unpaired t-test with Welch correction.

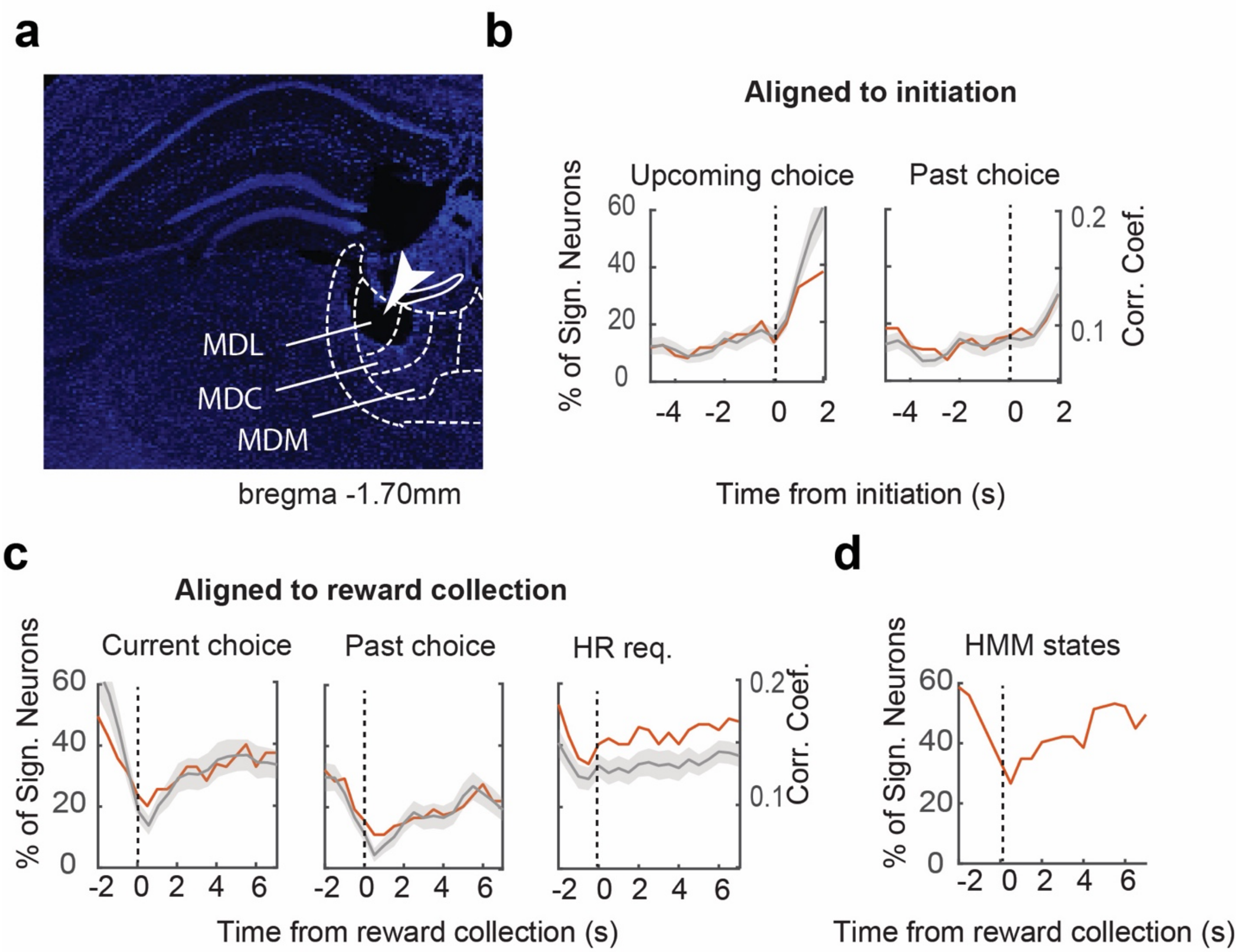


**Fig. S7 Neural activity in MD encodes HR required presses from reward collection to the initiation of the next trial, but shows higher choice encoding before reward collection and after initiation.**

**a,** Histology showing example lesion in the MD resulting from passing current through tetrodes before euthanasia. The white arrow indicates the lesion by current injection and the tetrode trace. MDL, lateral MD. MDC, central MD. MDM, medial MD. **b,** Percentage of neurons of which firing rate (FR) significantly correlates with the choice of the upcoming trial (top) or the choice of the past trial (bottom) from MD and the correlation coefficient of all neurons from MD at each time point aligned to initiation (grey line: mean; grey shade: mean+/-SEM). **c,** Percentage of neurons of which FR significantly correlates with the choice of the upcoming trial (left), the choice of the past trial (middle), or the HR required presses (right) from MD and the correlation coefficient of all neurons from MD at each time point aligned to Reward collection (grey line: mean; grey shade: mean +/-SEM). Note that the encoding of the current choice increases after the reward collection, it could reflect the reward volume difference associated with the choice. **d,** Percentage of neurons of which the FR is significantly different at exploration state and committed state at each time point aligned to reward collection. Req, required press numbers.

### Method

#### Animals

All animal-related work was performed under the guidelines of Division of Comparative Medicine (DCM), with protocol (# 2203-000-312 of Feng laboratory and #1017-077-20 of Halassa laboratory) approved by Committee for Animal Care (CAC) of Massachusetts Institute of Technology and was consistent with the Guide for Care and Use of Laboratory Animals, National Research Council 1996 (institutional animal welfare assurance no. A-3125-01). Only aged-matched male mice were used for all behavioral experiments; all other tests included age-matched males and females in proportional contribution across groups.

All behavioral studies were carried out and analyzed with the experimenter blinded to genotype. For all assays, mice were habituated in the test facility for 1 hour prior to starting the task. Each cohort of mice was used for maximally three behavioral tests with at least 5 days' break between tasks.

#### Generation of Grin2a$^{Y700X+/-}$ mice

*Gene editing.* Grin2a$^{Y700X+/-}$ mice were generated by CRIPSR-mediated knock in in zygote from C57BL/6 strain using standard procedures. crRNAs were designed using the CRISPR Guide RNA Design Tool (https://www.benchling.com/crispr) and tested by SURVEYOR assay (Transgenomic) according to the manufacturer's protocol. Donor DNA was designed as long ssDNA with homology arms with about 200 bp of size on 5' side and 150 bp of size on 3' side to flank the desired point mutation site. Briefly, crRNA (Synthego) and tracrRNA (Synthego) were annealed and mixed with Cas9 protein (NEB) to a final concentration of 8.7ng/µl crRNA, 14.3ng/ul tracrRNA and 30 ng/µl Cas9 protein to form a ribonucleotide protein complex. The ssDNA(IDT) was added to the mix and the cocktail was microinjected into the zygotes then transferred to the pseudo pregnant recipients. Mice were genotyped by Transnetyx (Cordova, TN).

*Breeding.* Heterozygous c57/BL6 male mice were group housed with WT female c57/BL6 mice to breed Heterozygous c57/BL6 male mice. Mice used for all experiments come from cages where Female WT S129 mice was paired with heterozygous c57/BL6 male mice to ensure 50% mutant and 50% WT littermates, normal maternal care and, 1:1 mixed background. New litter was tattooed on the toes for identification at P10 and the tail tissues were collected on the same day. Mice were weaned at P21 and were group housed by sex and genotypes. Only same mice with the same genotype were housed in the same cage to avoid abnormal social interaction between mice of different genotypes.

#### Western blot

To validate the expression level of Grin2a, we conducted Western blotting using hemisphere samples from WT, het and KO group at P14. The selected brain region encompassed half of the brain, excluding the olfactory bulb (OB) and cerebellum. The Western blotting procedures were carried out following the protocol previously described[1]. Immunoblotting was performed using the following antibodies: NR2A (Millipore Sigma, 04-901, 1:1000) and GAPDH (Santa Cruz Biotech, sc-32233, 1:1000).

#### Behavior screening

All behavior assay experiments were conducted during the light cycle from 7am to 7pm. Male WT and heterozygous littermates between 2 and 4 months of age were used for each experiment. All animals undergoing the open field and zero maze tests were handled by the investigator conducting the experiment for five minutes per day for three consecutive days leading up to the experiment. Investigators were blinded to genotypes for the entire duration of data collection and analysis of all behavior experiments.

*Open field test.* Group-housed mice were first habituated to a dark room containing the open-field experimental setup for one hour prior to the start of the experiment. Each mouse was individually placed at the bottom left corner of a four-cornered Plexiglass chamber with dimensions 40 x 40 x 30 cm (WLH). Prior to the first experiment and after all subsequent experiments, the chambers were cleaned with 70% ethanol and then thoroughly dried. For 60 minutes, motor activity was captured through automated infrared photobeam sensors through the Omnitech Digiscan apparatus (AccuScan Instruments, Columbus, OH). Spontaneous locomotion was measured by the total distance traveled and anxiety-like behavior was assessed using the times spent in the center versus the periphery of the arena.

*Elevated O-Maze.* Group-housed mice were habituated to the O-maze experiment room for one hour prior to the start of the experiment. The O-maze test apparatus consists of an oval-shaped arena elevated approximately 2.5 feet above the floor and is divided into 2 open-arm chambers and 2-closed arm chambers. Indirect lighting was adjusted such that the open-arm chambers were between 50-60 lux and the closed-arm chambers were between 5-10 lux. Each animal was placed at the border of an open and closed arm section with its head positioned towards the closed arm. Each mouse was given 5 minutes to explore the apparatus. Data collection of total time spent in open versus closed arms was scored automatically using the Noldus Observer software; anxiety-like behaviors were evaluated using the proportion of time spent in open-arm versus closed arm areas (Leesburg, VA).

*Sucrose preference test*. Sucrose preference test was conducted in each mouse's home cage. Prior to the experiment, the mice were single-housed and habituated to two identical spout bottles containing water. Following 72 hours of habituation, the two identical bottles were replaced with one bottle containing water and the other containing 2% sucrose solution. The total length of the experiment lasted 24 hours. At the 12-hour mark, the locations of the spout bottles were reversed to avoid any bias due to a preference for a specific side. The bottles were weighed at the beginning and end of the 24-hour period to measure sucrose solution and water consumed. Percent sucrose consumed was defined as the amount of sucrose solution consumed over the total liquid (water and sucrose solution) consumed.

*Prepulse inhibition of startle response test*. Startle response test was conducted in the set up from SR-Lab system.Prior to the experiment, the mice were habituated to the startle response chamber for 10 mins everyday for consecutively days until the mice didn't urinate or poop in the chamber. 24 hours following the habituation was finished, each mice was presented with 20 105db tone pulses (100ms) in the background of 60db white noise with randomized interval from 15 seconds to 25 seconds. And then mice were presented with three types of stimulus: 105db tone pulses, 85db tone pulses, and 85+105 pulses. Each type of stimulus was presented with the mice for 10 times with randomized order. Intervals between trials was randomized between 15 seconds to 25 seconds. Startle responses were recorded for 100ms after each tone was terminated. Startle response amplitude were analyzed by Matlab. Startle response (SR) amplitude was calculated as the peak within 50ms after each 105db tone. Prepulse inhibition ratio was calculated as (SR(105db)-SR(85+105db) )/SR(105db).

#### Lever-pressing task

Behavioral setup. Behavioral training and testing took place in gridded floor-mounted, custom-built chamber made of sheet metal covered with a thin layer of antistatic coating for electrical insulation (dimensions in cm: length, 15.2; width, 12.7; height, 24). Each chamber was equipped with an initiation port, two levers on each side, and one reward collection port in the center of the front wall. The initiation port, levers, and reward collection ports were 3-D printed, and all equipped with an IR LED/IR phototransistor pair (Digikey) for either nose poke detection or counting pressing numbers. The initiation port was located in the center of the chamber, about 6cm away from the front of the wall. Two plastic levers were mounted on the sides of the chamber, one on each side. One reward collection port was mounted in the center of the front wall with a syringe barrel mounted. The syringe barrel was connected to a single syringe pump (New Era Pump Systems) with silicone tubing. Evaporated milk was used as the reward. The ports and levers were controlled by an Arduino Mega microcontroller (Ivrea) for trial logic control. An infrared camera (Logitech) was attached to the top of the inside of the box to allow observation during sessions.
Training. Prior to training, all mice were food-restricted and maintained at 85–90% of their *ad libitum* body weight. Training occurred in 5 phases and took about 1-2 weeks. For all stages, mice were moved to the next phase of training if they performed more than 50 successful trials within 40 minutes. First, mice were placed into the testing chamber to acclimate. In this stage, it required the mice to press randomly assigned left or right lever one time to collect one reward (3ul milk) for building press-reward association. Second, the required press number will be increased to 3. Third, the required press number will be increased to 6 for the mice to collect the reward. Fourth, it required the mice to poke the initiation port for 50ms and then the randomly assigned lever would be enabled. Then the mice need to press the lever to collect the reward as during phase one. Usually, we would add a drop of milk at the initiation poke for the first few trials to guide the mice to poke. This stage is for training the mice to gain the 'initiation'--'pressing lever'--'collect reward' sequence. Fifth, after the mice were able to build this sequence, we increased the required press number to 6. After the mice fulfilled the training requirement in this stage, they were moved to the behavioral testing phase.

Testing. During the behavioral testing phase, one side of the lever was randomly assigned as the high reward (HR), and the other side was the low reward (LR) side. Fulfilling the HR lever pressing requirement will lead to a high reward: 9ul of milk while the LR lever leads to 3ul of milk. The initial requirement for the HR lever is 1 press while the requirement for the LR lever is 6 presses. After mice were put in the behavioral chamber, mice usually would randomly sample HR and LR lever and gradually commit to the HR side. After the mice showed preference for the HR lever (consecutively for 4 times), we started to increase the requirement to collect rewards on HR choices by one press every time the mice successfully finished a HR trial. If the animal chooses LR or didn't successfully fulfill the HR required press numbers in the current trial, the requirement for HR choice doesn't change for the next trial. If the mice didn't finish the required pressing numbers on the initially chosen side and pressed the other side, the trail would be 'aborted', and the mice would have a 30s time-out punishment. If the mice didn't finish the required pressing numbers and paused pressing for more than 10 s, the trial will be ended and considered 'incomplete'. If the mice choose the LR lever and get rewarded on LR choices consecutively 6 times, we consider the mice fully committed to the LR side, and a block is finished. After finishing one block, the requirement for HR was reset to one press and a new block started. If the mice didn't commit to the LR lever within 200 trials, a block was also terminated, and a new block will start.

_Data Analysis._ For each block, the probably of choosing HR (p(HR)) at each HR required press number was calculated by the proportion of HR choices of the trial numbers at each given HR required press. All blocks were cut off at HR required presses at 50. If mice committed to LR side before HR required presses at 50, the p(HR) at HR required presses higher than the committed HR requirement until 50 was considered 0. Block start was calculated as the after the mice consecutively get rewarded on HR lever for 4 times. Block end was calculated as after the mice consecutively get rewarded on LR lever for 6 times. Block length was calculated as from block start to block end. Optimality was calculated as the ratio of the reward the animal can get of each press to the reward of each press of the optimal policy.

_Optogenetic inhibition._ For laser blocks, laser pulses of blue (473 nm for iC++ activation) light at an intensity of 5-7 mW (measured at the tip of the optic fibers) were delivered pseudo-randomly on 50% of the blocks. During optogenetic experiments, laser stimulation occurred after the mice collected the reward of each trial and the light was terminated when the animal initiated the next trial. Light on and off blocks were randomized.

_Optogenetic activation._ For stabilized step function opsin (SSFO, hChR2(C128S/D156A)) experiments a 50-ms pulse of blue (473 nm, 5-7 mW intensity) light at the beginning of the session was delivered to activate the opsins and a 50-ms pulse of red (603 nm, 8 mW intensity) light to terminate activation at the end of the session. An Omicron-Laserage lighthub system (Dudenhofen) was used for all optogenetic manipulations. Each session is one hour and up to one session/day. Light on and off sessions were randomized.

**Virus injection and optic-fiber implantation**

Injections were performed using a quintessential stereotactic injector (QSI, Stoelting). Mice were deeply anaesthetized using 1% isoflurane. All viruses were obtained through UNC Chapel Hill, virus-vector core or addgene. Bilateral injections of AAV1-camkII-iC++-mcherry (400 nl) were used for the mediodorsal thalamus. For SSFO experiments, AAV-CamKIIa-SSFO-mcherry was injected bilaterally into the mediodorsal thalamus (400 nl). Optical fiber was implanted 0.2mm above the virus injection site bilaterally. Following virus injection and fiber implantation, animals were allowed to recover for at least two weeks for virus expression to take place before the start of behavioral testing or tissue collection.

**Multi-electrode array construction and implantation**

Custom-built multi-electrode drives targeting PFC and MD were built as previously described[2,3]. Custom multi-electrode array scaffolds (drive bodies) were designed using 3D CAD software (SolidWorks) and printed in Accura 55 plastic (American Precision Prototyping) as described previously. Prior to implantation, each array scaffold was loaded with 12–18 independently movable microdrives carrying 12.5-μm nichrome (California Fine Wire Company) stereotrodes or tetrodes. Electrodes were pinned to custom-designed, 96-channel electrode interface boards (EIB, Sunstone Circuits) along with a common reference wire (A-M systems). During implantation, mice were deeply anaesthetized with 1% isoflurane and mounted on a stereotaxic frame. A craniotomy was drilled centered at AP −3 mm, ML 2.5 mm for V1 (1.5 × 1.5 mm) or at AP −1 mm, ML 1.2 mm for MD recordings (approximately 2 × 2 mm). The dura was carefully removed, and the drive implant was lowered into the craniotomy using a stereotaxic arm until stereotrode tips touched the cortical surface. Surgilube (Savage Laboratories) was applied around electrodes to guard against fixation through dental cement. Stainless-steel screws were implanted into the skull to provide electrical and mechanical stability and the entire array was secured to the skull using dental cement.

### Electrophysiological recordings

Signals from tetrodes were acquired using a Neuralynx multiplexing digital recording system (Neuralynx) through a combination of 32- and 64-channel digital multiplexing headstages plugged into the 96-channel EIB of the implant. Signals from each electrode were amplified, filtered between 0.1 Hz and 9 kHz and digitized at 30 kHz. For thalamic recordings, tetrodes were lowered from the cortex into the mediodorsal thalamus over the course of 1–2 weeks where recording depths ranged from −2.8 to −3.2 mm DV. Following acquisition, spike sorting was performed offline on the basis of the relative spike amplitude and energy within electrode pairs using the MClust toolbox (https://github.com/adredish/MClust-Spike-Sorting-Toolbox) .[4]

### Electrophysiology Data analysis:

*Correlation:* 273 units from MD were recorded across 12 selected sessions (Only sessions in which animals have performed more than 100 trials and committed to LR before HR cost reached 60 and were included). Neurons that did not fire for the entire 7 seconds of correlation and decoding window in any trials and the median firing rate lower than 1 Hz during the intertrial interval were excluded. For prefrontal cortex, neurons were clustered based on the waveform and firing rate to separate fast spiking neurons from regular spiking neurons, and only regular spiking neurons were included. Therefore, only 109 MD spiking units were included for the analysis. The firing rate was calculated at a 2-second sliding window with 0.5 second sliding steps, and linear correlation was performed between firing rate at each time point to target value (cost value). In HMM states, instead of linear regression, we perform one-way ANOVA to test if firing rate is different between states.

*Decoding:* The decoding methods followed the methods previously described[5] , using Support Vector Machine (SVM) implemented through LIBSVM and the Matlab Neural Decoding Toolbox developed by Meyers[6]. 273 neurons were recorded from MD across 12 selected sessions (Only sessions in which animals have performed more than 30 trials and committed to LR before HR cost reached 60 and were included). Neurons that did not fire for the entire 7 seconds of correlation and decoding window in more than 10% of trials and median firing rate lower than 1 Hz during the intertrial interval were excluded. The firing rate of each neuron was calculated for each 600ms sliding window with 200ms overlapped between each sliding window. In decoding reward value, trials were divided into 2 categories, HR cost is larger and equal to 18 vs HR cost is smaller than 18. In decoding hmm states, trials were also divided into 2 categories, exploration state vs commitment state.

### Behavior Data Analysis

*HMM model:* We used the Hidden Markov Model to determine the states within each block objectively. The emission rate of each state is pre-determined as follows: HR-preferred state is defined as 80% probability of choosing HR; exploration state is defined as 50% probability of choosing HR and LR; LR-preferred state is defined as 80% probability of choosing LR. We first fitted the model to action data of each animal to determine the transition probability between states. Optimization of the model is done by finding the most likely transition probability for the given action sequence and predetermined emission rate using MATLAB function 'hmmdecode' and 'fmincon' to minimize negative log likelihood to identify the best fitting transition probabilities. Then we used the fit transition probabilities to categorize each trial as 1 of the 3 states based on both the emission rate (probability of choosing HR) and the transition rate (probability of staying at the same state vs transition to other states) using MATLAB function 'hmmdecode'. From this, we get length (number of trials) of each state for each block for each animal, and the bar graph was plotted using average across blocks of animals of the same transgenic group (Mutant vs WT).

*Bayesian model:* In brief, Bayesian inference was used to model how an animal performs value-based decision-making using belief updating. In this model, we assume an animal's choice is based on the relative inferred value of two options, HR and LR. The inferred value for each option is then defined as the ratio between reward size and cost (number of presses required), thus putting values in the quantity of reward achieved per lever press. We model the cost of HR and cost of LR as a t distribution with fixed shape, and we assume the reward value and the mean of LR cost are known for certain. Since the reward size is fixed for each option (HR/LR) and LR cost is fixed, the model dynamics rest on inferences about the cost (number of presses required) of the HR option. On each trial, the model uses probability distributions over the inferred value of HR and LR to compute the probability that HR is greater than LR, and then selects action HR according to this probability and LR otherwise.

Free parameters in the model are: α (belief updating rate, determining the relative weighting between prior and evidence), $\sigma_{HR}$ (t=0) (initial uncertainty associated with the HR cost distribution), $\sigma_{LR}$ (uncertainty of LR cost distribution), and $\delta$ (perceptual noise parameter).

The inferred value of the HR option was updated each time an animal completed a trial and thus received new evidence about the current level of effort required (E, new information of cost). The model assumed that evidence is corrupted by perceptual noise $\delta$ , such that new evidence is drawn from a uniform distribution ranging from $E - \delta \cdot \mathrm{E}$ to $E + \delta \cdot \mathrm{E}$. The update rule for updating parameters of the distribution of inferred HR cost ($\mu_{HR}$ *and* $\sigma_{HR}$) is based on Bayesian inference assuming unknown mean and standard deviation of the prior distribution derived by Murphy [7], where updating rule of $\mu_{HR}$ is the same as in reinforcement learning $\mu_{HR}\ (t+1) = (1-\alpha) \cdot \mu_{HR}\ (t) + \alpha \cdot \hat{E}$.

Action selection is then performed by drawing 200 samples from HR and LR cost distributions, which were assumed to be t-distributions with shape parameter equal to 1:

$$HR\_cost \ = t\ location\ scale\ distribution\ (\mu = \mu_{HR}, \sigma = \ \sigma_{HR}, \nu = 1)$$
$$LR\_cost \ = t\ location\ scale\ distribution\ (\mu = \mu_{LR}\ , \sigma = \sigma_{LR}, \nu = 1)$$

Note that while parameters of the HR distribution were updated dynamically, those of the LR were fixed, with $\mu_{LR}$ being fixed to the actual effort required for the low cost option (6) and $\sigma_{LR}$ fit as a free parameter.

The model was fit by maximizing likelihood of animal choice data aggregated across animals and sessions for each group. Parameters were estimated using a grid search over a range of parameter values: $\alpha$: $[0.1, 0.2, \dots\ to\ 0.9\ with\ 0.1\ increment]$, $\delta$: $[0.5, 1, ..4\ with\ 0.5\ increment]$, $\sigma_{HR}$: $[0.1, 0.2, 0.3 \dots 0.8\ with\ 0.1\ increment]$, $\sigma_{LR}$: $[1, 2, 3, 4, 5]$. Parameters were fitted to behavioral data from each individual animal and parameters fitted to animals in each group were compared (Fig. S6c).

Simulations (Fig S6b) were performed by endowing the model with the best-fitting parameter values for data pooled from all animals in each group and allowing the model to generate its own action on each trial, rather than report the likelihood of the animal action.

**Functional ultrasound imaging**

Surgery. Functional ultrasound imaging experiments were performed on mice aged 3-4 months as described[8]. Mice were deeply anaesthetized using 1% isoflurane and were head fixed on a quintessential stereotactic frame. Skull bones were removed from AP -2.7mm to 2.5mm and lateral -2mm to 2mm and replaced with ultrasound compatible plastic polymer. The plastic polymer was fixed by dental cement without blocking the imaging window. Silicone rubber was applied on top of the imaging window to provide protection. Mice were allowed to recover from the surgery for a week before the imaging session.

Imaging. During the imaging session, mice were head fixed and anaesthetized using 1.2% isoflurane to make sure that the mice keep stable anaesthetized state during the scanning session. Heart rate and oxygen saturation were monitored during the scanning session. Protective silicone rubber was removed, and ultrasound gel was applied to make sure there's no bubble between the scanning detector and the skull. The ultrasound detector was placed sagittal to the brain and moved from left to right during the scanning. Each animal was scanned for 30 minutes.

**Functional Ultrasound Data Analysis**

Baseline processing. Pre-processing steps roughly follow the steps in fMRI processing[9]. Functional ultrasound data was first manually aligned to the brain atlas in Icostudio (Iconeus, Paris, France). The atlas alignment matrix (rotation and transition matrix) and raw data were output to .h5 file and processed in MATLAB (Mathwork, Natick, MA, USA). We only used signals between 800 to 1400 seconds after putting on the nose cone for isoflurane anesthesia. We selected the window since we found the signals dropped gradually and became stable before 800th second and was stable without large baseline fluctuation. We first acquired a synchronized volumetric signal through **interpolating** signals of frames scanned at different time points at the same specified time point (sampling frequency of interpolation = (# of frames)*(volumetric-scanning-rate)). Then we **filtered (temporal domain)** the signal from each voxel with a non-shifted bandpass butterworth filter between 0.002 Hz to 0.12 Hz with filter order equal to 4 (filtfilt function). Then we **despiked** the signal from each voxel through removing the 2-second window before and after the time point where the extreme value happened. The extreme value threshold was defined by signals exceeding 2-fold of standard deviation of the overall signals from that voxel. The signal was then downsample (temporally) to original volumetric scanning frequency to reduce the size of data. The volumetric signal was then **rotated** and **transited** to align to the common atlas using the rotation and transition matrix output from Icostudio. The rotated volumetric data from each animal was then **resampled** at 0.03 mm x 0.03 mm x 0.03 mm spacing to get aligned volumetric data across animals. Then the data is spatially filtered with a 3-dimensional Gaussian filter at 2 pixel x 2 pixel x 2 pixel window (MATLAB function imgaussfilt3 (volumetric_data_at_one_time_point, 2).

*Local connectivity.* Local connectivity is calculated by averaging the cross-correlation of the 8 surrounding voxels at the same scanning frame (same sagittal frame with the same mediolateral coordinates).

*Significant cluster selection.* The selection of clusters of interest followed the methods previous described[10] and discussion with William Menegas and Frederico Azevedo. We first calculated the Cohen's d value between WT and mutant animal groups for each voxel. Then we thresholded the voxels with Cohen's d equal to and larger than 0.6 to be 1s, and rest to be 0s. Connected regions were identified from the thresholded volume. The cluster of volumes with volume size larger than 425 pixels was selected as regions of interest. The threshold of volume size was selected based on the p value from the permutation test described below.

*Significant test.* To select only the significant clusters, we ran a permutation test to calculate the chances of forming the cluster size from a set of randomized shuffled samples. We pooled all the pixel-wise time series data from all WT and mutant animals' data (19 animals total). For each round of the test, we randomly selected some pixel-wise time series from this pool to form a volume of scanning size for each sample to form 10 samples for group A and 9 samples for group B. We then processed the shuffled samples using the exact same temporal and spatial filtering methods, local connectivity calculation, group-wise comparison and clustering methods as we used for the actual samples. In the end of the round, we measured the largest cluster size showing the difference between group A and B with our predefined criteria for the actual samples. After 500 rounds, we got 500 largest cluster size numbers and plotted the distribution of the cluster size to calculate the possibility of forming each cluster size. We picked 425 pixels as the pixel threshold where the chances of forming cluster size above this number is less than 0.01 ($P<0.01$).

Displayed. The ROIs was then dilated with a sphere of radius of 3 pixels, followed by eroding with a sphere of radius of 5 pixels, dilating with a sphere of radius of 3 pixels and ended by eroding with a sphere of radius of 3 pixels for noise removal for image display purpose.

### Histology

Histology was performed for the tetrodes position and virus expression area. For histological verification of electrode position, drive-implanted mice were lightly anesthetized using isoflorane and small electrolytic lesions were generated by passing current (10 µA for 20 s) through the electrodes. All mice were then deeply anesthetized and transcardially perfused using phosphate-buffered saline (PBS) followed by 4% paraformaldehyde. Brains were dissected and postfixed overnight at 4 °C. Brain sections (50 µm) were cut using a vibratome (LEICA) and fluorescent images were obtained on a confocal microscope (LSM800, Zeiss). Confocal images are shown as maximal projection of 10 confocal planes, 20 µm thick.

### Statistical Testing

For all experiments except for the functional ultrasound studies, no statistical tests were done to determine the sample size. Required mouse numbers were assumed based on previous publications with similar approaches[1,11]. For functional ultrasound studies, the number of animals per group was pre-defined by power analysis from preliminary data. Animal identity was blinded to the data analyst and genotypes were revealed after the results were revealed. Data were first tested for normality using the Shapiro–Wilk test. For normally distributed data, Unpaired t test was used to compare between 2 groups and 1-way ANOVA was used to compare multiple groups.

For experiments involving multiple conditions (optogenetic manipulation, genotypes), two-way ANOVA with Tukey's multiple comparison test was used for multiple comparisons. T-test with multiple comparison correction (Welch correction) was used for the functional ultrasound experiment. For data set that didn't pass the normality test, the Kolmogorov-Simirnov D test was used to compare between 2 groups.


1. Zhou, Y. *et al.* Mice with Shank3 Mutations Associated with ASD and Schizophrenia Display Both Shared and Distinct Defects. *Neuron* **89**, 147–162 (2016).
2. Liang, L. *et al.* Scalable, Lightweight, Integrated and Quick-to-Assemble (SLIQ) Hyperdrives for Functional Circuit Dissection. *Front. Neural Circuits* **11**, 8 (2017).
3. Brunetti, P. M. *et al.* Design and fabrication of ultralight weight, adjustable multi-electrode probes for electrophysiological recordings in mice. *J. Vis. Exp. JoVE* e51675 (2014) doi:10.3791/51675.
4. David Redish. MClust, Spike sorting toolbox.
5. Rikhye, R. V., Gilra, A. & Halassa, M. M. Thalamic regulation of switching between cortical representations enables cognitive flexibility. *Nat. Neurosci.* **21**, 1753–1763 (2018).
6. Meyers, E. M. The neural decoding toolbox. *Front. Neuroinformatics* **7**, 8 (2013).
7. Murphy, K. Conjugate Bayesian analysis of the Gaussian distribution. (2007).
8. Brunner, C. *et al.* Whole-brain functional ultrasound imaging in awake head-fixed mice. *Nat. Protoc.* **16**, 3547–3571 (2021).
9. Rashid, B. *et al.* Classification of schizophrenia and bipolar patients using static and dynamic resting-state fMRI brain connectivity. *NeuroImage* **134**, 645–657 (2016).
10. Zhou, Y. *et al.* Atypical behaviour and connectivity in SHANK3-mutant macaques. *Nature* **570**, 326–331 (2019).
11. Schmitt, L. I. *et al.* Thalamic amplification of cortical connectivity sustains attentional control. *Nature* **545**, 219–223 (2017).
